## Supplementary material for "Correlative SHG-AFM imaging workflow for label-free quantitative analysis of collagen structure–function relationships": LaTEX: paperpile.bib

@ARTICLE{Jeong2018-rf,
  title     = "Visualization of drug distribution of a topical minocycline gel
               in human facial skin",
  author    = "Jeong, S and Hermsmeier, M and Osseiran, S and Yamamoto, A and
               Nagavarapu, U and Chan, K F and Evans, C L",
  journal   = "Biomedical optics express",
  publisher = "opg.optica.org",
  volume    =  9,
  number    =  7,
  pages     = "3434--3448",
  year      =  2018
}

@ARTICLE{Hwang2025-ik,
  title     = "A robust method for autofluorescence-free immunofluorescence
               using high-speed fluorescence lifetime imaging microscopy",
  author    = "Hwang, Wonsang and McPartland, Tyler and Jeong, Sinyoung and
               Evans, Conor L",
  journal   = "Sci. Rep.",
  publisher = "Springer Science and Business Media LLC",
  volume    =  15,
  number    =  1,
  pages     =  5503,
  abstract  = "Autofluorescence from endogenous biomolecules presents a
               significant challenge in immunofluorescence microscopy, often
               severely hindering the detection of specific fluorescence
               signals. While various techniques, including chemical- and
               quencher-based methods, photobleaching, and digital-based
               protocols like autofluorescence image subtraction, have been
               developed to suppress autofluorescence, each approach has
               inherent limitations that restrict their efficacy across
               different tissue types. Fluorescence lifetime imaging microscopy
               (FLIM) offers a powerful alternative by leveraging the distinct
               lifetime-spectrum profiles of fluorophores to differentiate
               specific immunofluorescence signals from autofluorescence.
               However, traditional FLIM methods are slow and have hampered its
               use for autofluorescence suppression in routine imaging
               applications. In this study, we demonstrate that GPU-accelerated
               high-speed FLIM can effectively separate autofluorescence from
               immunofluorescence signals in various tissue samples, achieving
               high throughput to meet the demands of biomedical and clinical
               workflows. Furthermore, our findings show that the FLIM-based
               autofluorescence suppression method enhances the correlation of
               immunofluorescence images with immunohistochemistry data,
               outperforming other methods like chemically-assisted
               photobleaching and hyperspectral imaging. These results highlight
               the potential of the high speed FLIM to improve the reliability
               of immunofluorescence microscopy significantly.",
  month     =  feb,
  year      =  2025,
  keywords  = "Autofluorescence; Fluorescence lifetime; Immunofluorescence",
  language  = "en"
}

@ARTICLE{de-Palma2004-by,
  title     = "Immunohistochemistry of the enthesis organ of the human Achilles
               tendon",
  author    = "de Palma, Luigi and Marinelli, Mario and Memè, Luca and Pavan,
               Matteo",
  journal   = "Foot Ankle Int.",
  publisher = "SAGE Publications",
  volume    =  25,
  number    =  6,
  pages     = "414--418",
  abstract  = "BACKGROUND: The insertional region of the Achilles tendon
               includes: the fibrocartilage of the enthesis at the tendon-bone
               junction, the sesamoid fibrocartilage in the deep surface of the
               tendon, and the periosteal fibrocartilage covering the superior
               tuberosity of the calcaneus. The mechanism by which compressive
               stress and low oxygen concentration modulate the phenotypical
               expression of cartilaginous cells in mesenchymal tissue is still
               unknown. Tenascin-C isoform, with its anti-adhesive properties,
               has been hypothesized to have a role in cellular adaptation to
               compression. The purpose of this study was to define the
               immunohistochemical localization of tenascin-C in the enthesis
               organ of the Achilles insertion in man. METHODS: Five specimens
               of Achilles tendon and its insertion were obtained from adult
               patients during below-knee amputation surgery. To determine the
               histochemistry, for light microscopy, specimens were stained with
               hematoxylin-eosin, safranin-O, Verhoeff, alcian blue methods. For
               immunohistochemistry, the following antibodies were used:
               anti-S-100 protein; anti-chondroitin sulfate; anti-types I and II
               collagen, anti-type IV collagen, anti-type X collagen, and
               anti-tenascin-C. RESULTS: The enthesis, sesamoid, and periosteal
               fibrocartilages have been defined by a separate region of
               immunohistochemical labeling for S-100 antibody and type II
               collagen and chondroitin-sulfate. The sesamoid and periosteal
               fibrocartilages stained for tenascin-C, which in cartilaginous
               areas was closely associated with rounded cells in the
               extracellular matrix. CONCLUSION: By showing a relationship
               between chondrocyte-like cells and immunolabeling for tenascin-C
               in periosteal and sesamoid fibrocartilage, the present data
               suggest a role in the dynamic equilibrium in the formation of
               fibrocartilaginous foci in tendon areas exposed to compressive
               stress. CLINICAL RELEVANCE: The data support the surgical
               practice of the resection of the diseased tendinous portion in
               case of chronic Achilles tendinitis. The decompression removed
               the enthesis organ that in an inflammatory environment could be
               altered and would not contribute to the distribution of the
               tensive load. After the decompression, it was hypothesized that
               in tendon areas exposed to compressive stress there was
               restoration of the fibrocartilaginous foci that constitute the
               enthesis organ.",
  month     =  jun,
  year      =  2004,
  language  = "en"
}

@ARTICLE{Silva2022-dy,
  title     = "Enthesis: not the same in each localisation-a molecular,
               histological and biomechanical study",
  author    = "Silva, C P and Müller, S A and Quirk, N and la Vega R. E., De and
               Coenen, M J and Evans, C H and {Van Griensven M.}",
  journal   = "Eur. Cells Mater",
  publisher = "cris.maastrichtuniversity.nl",
  volume    =  44,
  pages     = "43--55",
  year      =  2022
}

@ARTICLE{Angrisani2022-eh,
  title     = "Histology of tendon and enthesis—Suitable techniques for specific
               research questions",
  author    = "Angrisani, N and Willbold, E and Kampmann, A and Derksen, A and
               Reifenrath, J",
  journal   = "Eur. Cell Mater",
  publisher = "researchgate.net",
  volume    =  24,
  pages     = "228--251",
  year      =  2022
}

@ARTICLE{Kot2024-lo,
  title     = "Loss of the long form of {Plod2} phenocopies contractures of
               Bruck syndrome-osteogenesis imperfecta",
  author    = "Kot, Alexander and Chun, Cora and Martin, Jorge H and Wachtell,
               Davis and Hudson, David and Weis, Maryann and Marks, Haley and
               Srivastava, Siddharth and Eyre, David R and Duran, Ivan and
               Zieba, Jennifer and Krakow, Deborah",
  journal   = "J. Bone Miner. Res.",
  publisher = "Oxford University Press (OUP)",
  volume    =  39,
  number    =  9,
  pages     = "1240--1252",
  abstract  = "Bruck syndrome is an autosomal recessive form of osteogenesis
               imperfecta caused by biallelic variants in PLOD2 or FKBP10 and is
               characterized by joint contractures, bone fragility, short
               stature, and scoliosis. PLOD2 encodes LH2, which hydroxylates
               type I collagen telopeptide lysines, a critical step for collagen
               crosslinking. The Plod2 global knockout mouse model is limited by
               early embryonic lethality, and thus, the role of PLOD2 in
               skeletogenesis is not well understood. We generated a novel Plod2
               mouse line modeling a variant identified in two unrelated
               individuals with Bruck syndrome: PLOD2 c.1559dupC, predicting a
               frameshift and loss of the long isoform LH2b. In the mouse, the
               duplication led to loss of LH2b mRNA as well as significantly
               reduced total LH2 protein. This model, Plod2fs/fs, survived up to
               E18.5 although in non-Mendelian genotype frequencies. The
               homozygous frameshift model recapitulated the joint contractures
               seen in Bruck syndrome and had indications of absent type I
               collagen telopeptide lysine hydroxylation in bone. Genetically
               labeling tendons with Scleraxis-GFP in Plod2fs/fs mice revealed
               the loss of extensor tendons in the forelimb by E18.5, and
               developmental studies showed extensor tendons developed through
               E14.5 but were absent starting at E16.5. Second harmonic
               generation showed abnormal tendon type I collagen fiber
               organization, suggesting structurally abnormal tendons.
               Characterization of the skeleton by μCT and Raman spectroscopy
               showed normal bone mineralization levels. This work highlights
               the importance of properly crosslinked type I collagen in tendon
               and bone, providing a promising new mouse model to further our
               understanding of Bruck syndrome.",
  month     =  sep,
  year      =  2024,
  keywords  = "Bruck syndrome; LH2; PLOD2; bone; collagen crosslinking;
               hydroxylysine; mouse model; osteogenesis imperfecta; tendon",
  language  = "en"
}

@ARTICLE{Camy2022-gs,
  title     = "Effects of hindlimb unloading and subsequent reloading on the
               structure and mechanical properties of Achilles tendon-to-bone
               attachment",
  author    = "Camy, Claire and Brioche, Thomas and Senni, Karim and Bertaud,
               Alexandrine and Genovesio, Cécile and Lamy, Edouard and Fovet,
               Théo and Chopard, Angèle and Pithioux, Martine and Roffino,
               Sandrine",
  journal   = "FASEB J.",
  publisher = "Wiley",
  volume    =  36,
  number    =  10,
  pages     = "e22548",
  abstract  = "While muscle and bone adaptations to deconditioning have been
               widely described, few studies have focused on the tendon
               enthesis. Our study examined the effects of mechanical loading on
               the structure and mechanical properties of the Achilles tendon
               enthesis. We assessed the fibrocartilage surface area, the
               organization of collagen, the expression of collagen II, the
               presence of osteoclasts, and the tensile properties of the mouse
               enthesis both after 14 days of hindlimb suspension (HU) and after
               a subsequent 6 days of reloading. Although soleus atrophy was
               severe after HU, calcified fibrocartilage (CFc) was a little
               affected. In contrast, we observed a decrease in non-calcified
               fibrocartilage (UFc) surface area, collagen fiber
               disorganization, modification of morphological characteristics of
               the fibrocartilage cells, and altered collagen II distribution.
               Compared to the control group, restoring normal loads increased
               both UFc surface area and expression of collagen II, and led to a
               crimp pattern in collagen. Reloading induced an increase in CFc
               surface area, probably due to the mineralization front advancing
               toward the tendon. Functionally, unloading resulted in decreased
               enthesis stiffness and a shift in site of failure from the
               osteochondral interface to the bone, whereas 6 days of reloading
               restored the original elastic properties and site of failure. In
               the context of spaceflight, our results suggest that care must be
               taken when performing countermeasure exercises both during
               missions and during the return to Earth.",
  month     =  oct,
  year      =  2022,
  keywords  = "Achilles tendon enthesis; collagen II; fibrocartilage; hindlimb
               suspension; mechanical properties; reloading",
  language  = "en"
}

@ARTICLE{Smith2011-tb,
  title     = "Hamstring contractures in children with spastic cerebral palsy
               result from a stiffer extracellular matrix and increased in vivo
               sarcomere length: Passive mechanical properties of muscle
               contracture",
  author    = "Smith, Lucas R and Lee, Ki S and Ward, Samuel R and Chambers,
               Henry G and Lieber, Richard L",
  journal   = "J. Physiol.",
  publisher = "Wiley",
  volume    =  589,
  number    = "Pt 10",
  pages     = "2625--2639",
  abstract  = "Cerebral palsy (CP) results from an upper motoneuron (UMN)lesion
               in the developing brain. Secondary to the UMNl esion,which causes
               spasticity, is a pathological response by muscle - namely,
               contracture. However, the elements within muscle that increase
               passive mechanical stiffness, and therefore result in
               contracture, are unknown. Using hamstring muscle biopsies from
               pediatric patients with CP (n =33) and control (n =19) patients
               we investigated passive mechanical properties at the protein,
               cellular, tissue and architectural levels to identify the
               elements responsible for contracture. Titin isoform, the major
               load-bearing protein within muscle cells, was unaltered in CP.
               Correspondingly, the passive mechanics of individual muscle
               fibres were not altered. However, CP muscle bundles, which
               include fibres in their constituent ECM, were stiffer than
               control bundles. This corresponded to an increase in collagen
               content of CP muscles measured by hydroxyproline assay and
               observed using immunohistochemistry. In vivo sarcomere length of
               CP muscle measured during surgery was significantly longer than
               that predicted for control muscle. The combination of increased
               tissue stiffness and increased sarcomere length interact to
               increase stiffness greatly of the contracture tissue in vivo.
               These findings provide evidence that contracture formation is not
               the result of stiffening at the cellular level, but stiffening of
               the ECM with increased collagen and an increase of in vivo
               sarcomere length leading to higher passive stresses.",
  month     =  may,
  year      =  2011,
  language  = "en"
}

@ARTICLE{Pena2005-rq,
  title     = "Chiroptical effects in the second harmonic signal of collagens
               {I} and {IV}",
  author    = "Pena, Ana-Maria and Boulesteix, Thierry and Dartigalongue,
               Thibault and Schanne-Klein, Marie-Claire",
  journal   = "J. Am. Chem. Soc.",
  publisher = "American Chemical Society (ACS)",
  volume    =  127,
  number    =  29,
  pages     = "10314--10322",
  abstract  = "We performed polarization-resolved surface second harmonic
               generation (SHG) experiments on thin films of collagen I and IV
               molecules, as well as conventional CD measurements. We found that
               collagen IV presents little CD and no SHG optical activity,
               whereas collagen I exhibits large chiroptical effects involving
               both one-electron and excitonic coupling mechanisms. We estimated
               that these chiral components enhance the SHG signal from
               fibrillar collagen in biological tissues by typically a factor of
               2. By comparing the distinct behaviors of collagens I and IV in
               SHG microscopy and in surface SHG experiments, we concluded that
               SHG microscopy is a sensitive probe of the micrometer-scale
               structural organization of collagen in biological tissues.",
  month     =  jul,
  year      =  2005,
  language  = "en"
}

@ARTICLE{Fuentes-Corona2019-dt,
  title     = "Second harmonic generation signal from type {I} collagen fibers
               grown in vitro",
  author    = "Fuentes-Corona, Cindy Grethel and Licea-Rodriguez, Jacob and
               Younger, Rebecca and Rangel-Rojo, Raul and Potma, Eric O and
               Rocha-Mendoza, Israel",
  journal   = "Biomed. Opt. Express",
  publisher = "Optica Publishing Group",
  volume    =  10,
  number    =  12,
  pages     = "6449--6461",
  abstract  = "We present a study of the optical second-order nonlinearity of
               type I collagen fibers grown in vitro via second harmonic
               generation (SHG) experiments and analyze the observed
               polarization-resolved SHG signal using previously reported SHG
               analytical expressions obtained for anisotropic tissue. Our
               results indicate that the effective second-order nonlinearity
               measured in the grown fibers is one order of magnitude lower than
               that of native collagen fibers. This is attributed to the
               formation of loose and dispersive fibrillar networks of thinner
               collagen fibrils that constitute the reassembled collagen fibers.
               This is confirmed by scanning electronic microscopy (SEM) imaging
               and the polarization dependence of the SHG signal. The measured
               values of the anisotropy parameter ρ of the reassembled collagen
               fibers are found to be similar to that obtained for native fibers
               on the relevant sub-µm scale.",
  month     =  dec,
  year      =  2019,
  language  = "en"
}

@ARTICLE{Johnson2021-ex,
  title     = "Superresolved polarization-enhanced second-harmonic generation
               for direct imaging of nanoscale changes in collagen architecture",
  author    = "Johnson, Peter B and Karvounis, Artemios and Singh, H Johnson and
               Brereton, Christopher J and Bourdakos, Konstantinos N and Lunn,
               Kerry and Roberts, James J W and Davies, Donna E and Muskens,
               Otto L and Jones, Mark G and Mahajan, Sumeet",
  journal   = "Optica",
  publisher = "Optica Publishing Group",
  volume    =  8,
  number    =  5,
  pages     = "674--685",
  abstract  = "Superresolution (SR) optical microscopy has allowed the
               investigation of many biological structures below the diffraction
               limit; however, most of the techniques are hampered by the need
               for fluorescent labels. Nonlinear label-free techniques such as
               second-harmonic generation (SHG) provide structurally specific
               contrast without the addition of exogenous labels, allowing
               observation of unperturbed biological systems. We use the
               photonic nanojet (PNJ) phenomena to achieve SR-SHG. A resolution
               of ∼ λ / 6 with respect to the fundamental wavelength, that is, a
               ∼ 2.3 -fold improvement over conventional or diffraction-limited
               SHG under the same imaging conditions is achieved. Crucially we
               find that the polarization properties of excitation are
               maintained in a PNJ. This is observed in experiment and
               simulations. This may have widespread implications to increase
               sensitivity by detection of polarization-resolved SHG by
               observing anisotropy in signals. These new, to the best of our
               knowledge, findings allowed us to visualize biological SHG-active
               structures such as collagen at an unprecedented and previously
               unresolvable spatial scale. Moreover, we demonstrate that the use
               of an array of self-assembled high-index spheres overcomes the
               issue of a limited field of view for such a method, allowing
               PNJ-assisted SR-SHG to be used over a large area. Dysregulation
               of collagen at the nanoscale occurs in many diseases and is an
               underlying cause in diseases such as lung fibrosis. Here we
               demonstrate that pSR-SHG allows unprecedented observation of
               changes at the nanoscale that are invisible by conventional
               diffraction-limited SHG imaging. The ability to nondestructively
               image SHG-active biological structures without labels at the
               nanoscale with a relatively simple optical method heralds the
               promise of a new tool to understand biological phenomena and
               drive drug discovery.",
  month     =  may,
  year      =  2021,
  language  = "en"
}

@ARTICLE{Ogura2019-ks,
  title     = "Texture analysis of second-harmonic-generation images for
               quantitative analysis of reticular dermal collagen fibre in vivo
               in human facial cheek skin",
  author    = "Ogura, Yuki and Tanaka, Yuji and Hase, Eiji and Yamashita,
               Toyonobu and Yasui, Takeshi",
  journal   = "Exp. Dermatol.",
  publisher = "Wiley",
  volume    =  28,
  number    =  8,
  pages     = "899--905",
  abstract  = "Second-harmonic-generation (SHG) microscopy is a powerful tool
               for in vivo visualisation of collagen fibres in human skin
               because of its specific collagen selectivity without the need for
               staining, non-invasiveness and high-resolution three-dimensional
               imaging. Although texture analysis of SHG images is a promising
               method for the quantitative analysis of well-orientated collagen
               fibre structure in the tendon and cornea, there are few attempts
               to assess cutaneous ageing. In this study, we applied two texture
               analysis techniques, namely autocorrelation (2D-AC) analysis and
               two-dimensional Fourier transform (2D-FT), to evaluate the
               age-dependent changes in reticular dermal collagen fibres in in
               vivo human cheek skin. Age-dependent changes in the reticular
               dermal collagen fibres of female subjects in their 20s, 40s and
               60s clearly appeared in these texture analyses. Furthermore, the
               parameter from 2D-AC analysis showed a significantly higher
               correlation with skin elasticity measured by a Cutometer® . These
               results clearly indicate that 2D-AC analysis of SHG images is
               highly promising for the quantitative evaluation of age-dependent
               change in facial collagen fibres as well as skin elasticity. An
               appropriate texture analysis will help to provide quantitative
               insight into collagen fibre structure and will be useful for the
               diagnosis of pathological conditions as well as cutaneous ageing
               in skin.",
  month     =  aug,
  year      =  2019,
  keywords  = "SHG; ageing; elasticity; non-invasive",
  language  = "en"
}

@ARTICLE{Aghigh2023-iu,
  title     = "Second harmonic generation microscopy: a powerful tool for
               bio-imaging",
  author    = "Aghigh, Arash and Bancelin, Stéphane and Rivard, Maxime and
               Pinsard, Maxime and Ibrahim, Heide and Légaré, François",
  journal   = "Biophys. Rev.",
  publisher = "Springer Science and Business Media LLC",
  volume    =  15,
  number    =  1,
  pages     = "43--70",
  abstract  = "Second harmonic generation (SHG) microscopy is an important
               optical imaging technique in a variety of applications. This
               article describes the history and physical principles of SHG
               microscopy and its more advanced variants, as well as their
               strengths and weaknesses in biomedical applications. It also
               provides an overview of SHG and advanced SHG imaging in
               neuroscience and microtubule imaging and how these methods can
               aid in understanding microtubule formation, structuration, and
               involvement in neuronal function. Finally, we offer a perspective
               on the future of these methods and how technological advancements
               can help make SHG microscopy a more widely adopted imaging
               technique.",
  month     =  feb,
  year      =  2023,
  keywords  = "Interferometry; Neuroimaging; Non-linear microscopy; Polarimetry;
               SHG",
  language  = "en"
}

@ARTICLE{Liu2023-at,
  title     = "Label-free, multi-parametric assessments of cell metabolism and
               matrix remodeling within human and early-stage murine
               osteoarthritic articular cartilage",
  author    = "Liu, Zhiyi and Hui Mingalone, Carrie K and Gnanatheepam, Einstein
               and Hollander, Judith M and Zhang, Yang and Meng, Jia and Zeng,
               Li and Georgakoudi, Irene",
  journal   = "Commun. Biol.",
  publisher = "Springer Science and Business Media LLC",
  volume    =  6,
  number    =  1,
  pages     =  405,
  abstract  = "Osteoarthritis (OA) is characterized by the progressive
               deterioration of articular cartilage, involving complicated
               cell-matrix interactions. Systematic investigations of dynamic
               cellular and matrix changes during OA progression are lacking. In
               this study, we use label-free two-photon excited fluorescence
               (TPEF) and second harmonic generation (SHG) imaging to assess
               cellular and extracellular matrix features of murine articular
               cartilage during several time points at early stages of OA
               development following destabilization of medial meniscus surgery.
               We detect significant changes in the organization of collagen
               fibers and crosslink-associated fluorescence of the superficial
               zone as early as one week following surgery. Such changes become
               significant within the deeper transitional and radial zones at
               later time-points, highlighting the importance of high spatial
               resolution. Cellular metabolic changes exhibit a highly dynamic
               behavior, and indicate metabolic reprogramming from enhanced
               oxidative phosphorylation to enhanced glycolysis or fatty acid
               oxidation over the ten-week observation period. The optical
               metabolic and matrix changes detected within this mouse model are
               consistent with differences identified in excised human cartilage
               specimens from OA and healthy cartilage specimens. Thus, our
               studies reveal important cell-matrix interactions at the onset of
               OA that may enable improved understanding of OA development and
               identification of new potential treatment targets.",
  month     =  apr,
  year      =  2023,
  language  = "en"
}

@ARTICLE{Finnoy2016-cd,
  title     = "Second harmonic generation imaging reveals a distinct
               organization of collagen fibrils in locations associated with
               cartilage growth",
  author    = "Finnøy, Andreas and Olstad, Kristin and Lilledahl, Magnus B",
  journal   = "Connect. Tissue Res.",
  publisher = "Informa UK Limited",
  volume    =  57,
  number    =  5,
  pages     = "374--387",
  abstract  = "PURPOSE: The articular-epiphyseal cartilage complex (AECC) is
               responsible for the expansion of the bone ends and serves the
               function of the articular cartilage in juvenile mammals. Bundles
               of collagen fibrils surrounding cells were in the literature
               observed more frequently near the articular surface of the AECC.
               The articular surface, the perichondrium, and cartilage canals
               are interfaces where appositional growth of the AECC has been
               demonstrated. The current study aimed to evaluate the potential
               of second harmonic generation (SHG) to locate the collagen fibril
               bundles near the articular surface and to examine whether a
               comparable collagen fibril organization could be observed near
               the other interfaces of the AECC. MATERIALS AND METHODS: The
               study included the femoral condyle of four piglets aged 82-141
               days. The forward and backward scattered SHG, and their ratio,
               was analyzed across the AECC using objectives with different
               numerical aperture. Two-photon-excited fluorescence was used to
               visualize cells. RESULTS: A similar pattern of collagen fibril
               organization was observed near the articular surface, around
               cartilage canals, and adjacent to the perichondrium. The pattern
               consisted of a higher ratio of forward to backward scattered SHG
               that increased relative to the surrounding matrix at lower
               numerical aperture. This was interpreted to reflect collagen
               fibril bundles in the territorial matrix of cells in these areas.
               CONCLUSIONS: The observed arrangement of collagen fibrils was
               suggested to be related to the presumed different growth activity
               in these areas and indicated that SHG may be used as an indirect
               and label-free marker for cartilage matrix growth.",
  month     =  sep,
  year      =  2016,
  keywords  = "Articular cartilage; cartilage canal; cartilage growth; collagen
               fibrils; second harmonic generation",
  language  = "en"
}

@ARTICLE{Crosignani2014-do,
  title     = "A deep tissue fluorescence imaging system with enhanced {SHG}
               detection capabilities: Deep Tissue Fluorescence Imaging System",
  author    = "Crosignani, Viera and Jahid, Sohail and Dvornikov, Alexander S
               and Gratton, Enrico",
  journal   = "Microsc. Res. Tech.",
  publisher = "Wiley",
  volume    =  77,
  number    =  5,
  pages     = "368--373",
  abstract  = "We describe a novel two-photon fluorescence microscopy system
               capable of producing high-quality second harmonic generation
               (SHG) images in thick turbid media by using an innovative
               detection system. This novel detection system is capable of
               detecting photons from a very large surface area. This system has
               proven effective in providing images of thick turbid samples,
               both biological and artificial. Due to its transmission detection
               geometry, the system is particularly suitable for detecting SHG
               signals, which are generally forward directed. In this article,
               we present comparative data acquired simultaneously on the same
               sample with the forward and epidetection schemes.",
  month     =  may,
  year      =  2014,
  keywords  = "deep tissue imaging; second harmonic generation; two-photon
               microscopy",
  language  = "en"
}

@ARTICLE{Hristu2021-vq,
  title     = "Influence of hematoxylin and eosin staining on the quantitative
               analysis of second harmonic generation imaging of fixed tissue
               sections",
  author    = "Hristu, Radu and Stanciu, Stefan G and Dumitru, Adrian and Paun,
               Bogdan and Floroiu, Iustin and Costache, Mariana and Stanciu,
               George A",
  journal   = "Biomed. Opt. Express",
  publisher = "Optica Publishing Group",
  volume    =  12,
  number    =  9,
  pages     = "5829--5843",
  abstract  = "Second harmonic generation (SHG) microscopy has emerged over the
               past two decades as a powerful tool for tissue characterization
               and diagnostics. Its main applications in medicine are related to
               mapping the collagen architecture of in-vivo, ex-vivo and fixed
               tissues based on endogenous contrast. In this work we present how
               H\&E staining of excised and fixed tissues influences the
               extraction and use of image parameters specific to
               polarization-resolved SHG (PSHG) microscopy, which are known to
               provide quantitative information on the collagen structure and
               organization. We employ a theoretical collagen model for fitting
               the experimental PSHG datasets to obtain the second order
               susceptibility tensor elements ratios and the fitting efficiency.
               Furthermore, the second harmonic intensity acquired under
               circular polarization is investigated. The evolution of these
               parameters in both forward- and backward-collected SHG are
               computed for both H\&E-stained and unstained tissue sections.
               Consistent modifications are observed between the two cases in
               terms of the fitting efficiency and the second harmonic
               intensity. This suggests that similar quantitative analysis
               workflows applied to PSHG images collected on stained and
               unstained tissues could yield different results, and hence affect
               the diagnostic accuracy.",
  month     =  sep,
  year      =  2021,
  language  = "en"
}

@ARTICLE{Roth1981-qi,
  title     = "Optical second-harmonic scattering in rat-tail tendon",
  author    = "Roth, S and Freund, I",
  journal   = "Biopolymers",
  publisher = "Wiley",
  volume    =  20,
  number    =  6,
  pages     = "1271--1290",
  abstract  = "AbstractThe angular dependence of optical second‐harmonic
               generation by native, wet, rat‐tail tendon is found to display a
               sharp, intense, forward peak superimposed on a broad background.
               The sharp peak is shown to imply long‐range polar order, while
               the broad background corresponds to that predicted for the random
               “up”/“down” array of collagen fibrils seen with the electron
               microscope. The degree of polar order is determined, and the
               dependence of the fibril diameter distribution on age and state
               of hydration is measured. The coherence length of tendon for
               harmonic generation and the absolute magnitude of the nonlinear
               susceptibility of the collagen fibril are also determined. The
               biological significance of these various findings is discussed.",
  month     =  jun,
  year      =  1981,
  language  = "en"
}

@ARTICLE{Liu2017-wm,
  title     = "Methods for quantifying fibrillar collagen alignment",
  author    = "Liu, Yuming and Keikhosravi, Adib and Mehta, Guneet S and Drifka,
               Cole R and Eliceiri, Kevin W",
  journal   = "Methods Mol. Biol.",
  publisher = "Springer New York",
  address   = "New York, NY",
  volume    =  1627,
  pages     = "429--451",
  abstract  = "Recent evidence has implicated collagen, particularly fibrillar
               collagen, in a number of diseases ranging from osteogenesis
               imperfecta and asthma to breast and ovarian cancer. A key
               property of collagen that has been correlated with disease has
               been the alignment of collagen fibers. Collagen can be visualized
               using a variety of imaging techniques including second-harmonic
               generation (SHG) microscopy, polarized light microscopy, and
               staining with dyes or antibodies. However, there exists a great
               need to easily and robustly quantify images from these modalities
               for individual fibers in specified regions of interest and with
               respect to relevant boundaries. Most currently available
               computational tools rely on calculation of pixel-wise orientation
               or global window-wise orientation that do not directly calculate
               or give visible fiber-wise information and do not provide
               relative orientation against boundaries. We describe and detail
               how to use a freely available, open-source MATLAB software
               framework that includes two separate but linked packages
               ``CurveAlign'' and ``CT-FIRE'' that can address this need by
               either directly extracting individual fibers using an improved
               fiber tracking algorithm or directly finding optimal
               representation of fiber edges using the curvelet transform. This
               curvelet-based framework allows the user to measure fiber
               alignment on a global, region of interest, and fiber basis.
               Additionally, users can measure fiber angle relative to manually
               or automatically segmented boundaries. This tool does not require
               prior experience of programming or image processing and can
               handle multiple files, enabling efficient quantification of
               collagen organization from biological datasets.",
  year      =  2017,
  keywords  = "Cancer imaging; Collagen alignment; Collagen organization;
               Collagen quantification; Curvelet transform; Extracellular
               matrix; Fiber tracking; Fibrillar collagen; Second-harmonic
               generation microscopy; Tumor-associated collagen signatures",
  language  = "en"
}

@ARTICLE{McLean2019-tb,
  title     = "High-speed collagen fiber modeling and orientation quantification
               for optical coherence tomography imaging",
  author    = "McLean, James P and Gan, Yu and Lye, Theresa H and Qu, Dovina and
               Lu, Helen H and Hendon, Christine P",
  journal   = "Opt. Express",
  publisher = "Optica Publishing Group",
  volume    =  27,
  number    =  10,
  pages     = "14457--14471",
  abstract  = "Quantifying collagen fiber architecture has clinical and
               scientific relevance across a variety of tissue types and adds
               functionality to otherwise largely qualitative imaging
               modalities. Optical coherence tomography (OCT) is uniquely suited
               for this task due to its ability to capture the collagen
               microstructure over larger fields of view than traditional
               microscopy. Existing image processing techniques for quantifying
               fiber architecture, while accurate and effective, are very slow
               for processing large datasets and tend to lack structural
               specificity. We describe here a computationally efficient method
               for quantifying and visualizing collagen fiber organization. The
               algorithm is demonstrated on swine atria, bovine anterior
               cruciate ligament, and human cervical tissue samples.
               Additionally, we show an improved performance for images with
               crimped fiber textures and low signal to noise when compared to
               similar methods.",
  month     =  may,
  year      =  2019,
  language  = "en"
}

@ARTICLE{Ranjit2015-ct,
  title     = "Imaging fibrosis and separating collagens using second harmonic
               generation and phasor approach to fluorescence lifetime imaging",
  author    = "Ranjit, Suman and Dvornikov, Alexander and Stakic, Milka and
               Hong, Suk-Hyun and Levi, Moshe and Evans, Ronald M and Gratton,
               Enrico",
  journal   = "Sci. Rep.",
  publisher = "Springer Science and Business Media LLC",
  volume    =  5,
  number    =  1,
  pages     =  13378,
  abstract  = "In this paper we have used second harmonic generation (SHG) and
               phasor approach to auto fluorescence lifetime imaging (FLIM) to
               obtain fingerprints of different collagens and then used these
               fingerprints to observe bone marrow fibrosis in the mouse femur.
               This is a label free approach towards fast automatable detection
               of fibrosis in tissue samples. FLIM has previously been used as a
               method of contrast in different tissues and in this paper phasor
               approach to FLIM is used to separate collagen I from collagen
               III, the markers of fibrosis, the largest groups of disorders
               that are often without any effective therapy. Often characterized
               by an increase in collagen content of the corresponding tissue,
               the samples are usually visualized by histochemical staining,
               which is pathologist dependent and cannot be automated.",
  month     =  aug,
  year      =  2015,
  language  = "en"
}

@ARTICLE{Chen2012-ac,
  title     = "Second harmonic generation microscopy for quantitative analysis
               of collagen fibrillar structure",
  author    = "Chen, Xiyi and Nadiarynkh, Oleg and Plotnikov, Sergey and
               Campagnola, Paul J",
  journal   = "Nat. Protoc.",
  publisher = "Springer Science and Business Media LLC",
  volume    =  7,
  number    =  4,
  pages     = "654--669",
  abstract  = "Second-harmonic generation (SHG) microscopy has emerged as a
               powerful modality for imaging fibrillar collagen in a diverse
               range of tissues. Because of its underlying physical origin, it
               is highly sensitive to the collagen fibril/fiber structure, and,
               importantly, to changes that occur in diseases such as cancer,
               fibrosis and connective tissue disorders. We discuss how SHG can
               be used to obtain more structural information on the assembly of
               collagen in tissues than is possible by other microscopy
               techniques. We first provide an overview of the state of the art
               and the physical background of SHG microscopy, and then describe
               the optical modifications that need to be made to a
               laser-scanning microscope to enable the measurements. Crucial
               aspects for biomedical applications are the capabilities and
               limitations of the different experimental configurations. We
               estimate that the setup and calibration of the SHG instrument
               from its component parts will require 2-4 weeks, depending on the
               level of the user's experience.",
  month     =  mar,
  year      =  2012,
  language  = "en"
}

@ARTICLE{Lim2019-lx,
  title     = "Harmonic generation microscopy 2.0: New tricks empowering
               intravital imaging for neuroscience",
  author    = "Lim, Hyungsik",
  journal   = "Front. Mol. Biosci.",
  publisher = "Frontiers Media SA",
  volume    =  6,
  pages     =  99,
  abstract  = "Optical harmonic generation, e.g., second- (SHG) and
               third-harmonic generation (THG), provides intrinsic contrasts for
               three-dimensional intravital microscopy. Contrary to two-photon
               excited fluorescence (TPEF), however, they have found relatively
               specialized applications, such as imaging collagenous and
               non-specific tissues, respectively. Here we review recent
               advances that broaden the capacity of SHG and THG for imaging the
               central nervous system in particular. The fundamental contrast
               mechanisms are reviewed as they encode novel information
               including molecular origin, spectroscopy, functional probes, and
               image analysis, which lay foundations for promising future
               applications in neuroscience.",
  month     =  oct,
  year      =  2019,
  keywords  = "intravital imaging; label-free imaging; non-linear optical
               microscopy; second-harmonic generation; third-harmonic generation",
  language  = "en"
}

@ARTICLE{James2021-zt,
  title     = "Recent advancements in optical harmonic generation microscopy:
               Applications and perspectives",
  author    = "James, Darian S and Campagnola, Paul J",
  journal   = "BME Front.",
  publisher = "American Association for the Advancement of Science (AAAS)",
  volume    =  2021,
  pages     =  3973857,
  abstract  = "Second harmonic generation (SHG) and third harmonic generation
               (THG) microscopies have emerged as powerful imaging modalities to
               examine structural properties of a wide range of biological
               tissues. Although SHG and THG arise from very different contrast
               mechanisms, the two are complimentary and can often be collected
               simultaneously using a modified multiphoton microscope. In this
               review, we discuss the needed instrumentation for these
               modalities as well as the underlying theoretical principles of
               SHG and THG in tissue and describe how these can be leveraged to
               extract unique structural information. We provide an overview of
               recent advances showing how SHG microscopy has been used to
               evaluate collagen alterations in the extracellular matrix and how
               this has been used to advance our knowledge of cancers, fibroses,
               and the cornea, as well as in tissue engineering applications.
               Specific examples using polarization-resolved approaches and
               machine learning algorithms are highlighted. Similarly, we review
               how THG has enabled developmental biology and skin cancer studies
               due to its sensitivity to changes in refractive index, which are
               ubiquitous in all cell and tissue assemblies. Lastly, we offer
               perspectives and outlooks on future directions of SHG and THG
               microscopies and present unresolved questions, especially in
               terms of overall miniaturization and the development of
               microendoscopy instrumentation.",
  month     =  jan,
  year      =  2021,
  language  = "en"
}

@ARTICLE{Williams2005-id,
  title     = "Interpreting second-harmonic generation images of collagen {I}
               fibrils",
  author    = "Williams, Rebecca M and Zipfel, Warren R and Webb, Watt W",
  journal   = "Biophys. J.",
  publisher = "Elsevier BV",
  volume    =  88,
  number    =  2,
  pages     = "1377--1386",
  abstract  = "Fibrillar collagen, being highly noncentrosymmetric, possesses a
               tremendous nonlinear susceptibility. As a result, second-harmonic
               generation (SHG) microscopy of collagen produces extremely bright
               and robust signals, providing an invaluable tool for imaging
               tissue structure with submicron resolution. Here we discuss
               fundamental principles governing SHG phase matching with the
               tightly focusing optics used in microscopy. Their application to
               collagen imaging yields several biophysical features
               characteristic of native collagen structure: SHG radiates from
               the shell of a collagen fibril, rather than from its bulk. This
               SHG shell may correspond to the supporting element of the fibril.
               Physiologically relevant changes in solution ionic strength alter
               the ratio of forward-to-backward propagating SHG, implying a
               resulting change in the SHG shell thickness. Fibrillogenesis can
               be resolved in immature tissue by directly imaging
               backward-propagating SHG. Such findings are crucial to the design
               and development of forthcoming diagnostic and research tools.",
  month     =  feb,
  year      =  2005,
  language  = "en"
}

@ARTICLE{Gillies2011-zi,
  title     = "Structure and function of the skeletal muscle extracellular
               matrix: Skeletal Muscle {ECM}",
  author    = "Gillies, Allison R and Lieber, Richard L",
  journal   = "Muscle Nerve",
  publisher = "Wiley",
  volume    =  44,
  number    =  3,
  pages     = "318--331",
  abstract  = "The skeletal muscle extracellular matrix (ECM) plays an important
               role in muscle fiber force transmission, maintenance, and repair.
               In both injured and diseased states, ECM adapts dramatically, a
               property that has clinical manifestations and alters muscle
               function. Here we review the structure, composition, and
               mechanical properties of skeletal muscle ECM; describe the cells
               that contribute to the maintenance of the ECM; and, finally,
               overview changes that occur with pathology. New scanning electron
               micrographs of ECM structure are also presented with hypotheses
               about ECM structure–function relationships. Detailed
               structure–function relationships of the ECM have yet to be
               defined and, as a result, we propose areas for future study.",
  month     =  sep,
  year      =  2011,
  keywords  = "Skeletal muscle; biomechanics; collagen; extracellular matrix;
               fibrosis",
  language  = "en"
}

@ARTICLE{Liu2020-uo,
  title     = "Fibrillar collagen quantification with curvelet transform based
               computational methods",
  author    = "Liu, Yuming and Keikhosravi, Adib and Pehlke, Carolyn A and
               Bredfeldt, Jeremy S and Dutson, Matthew and Liu, Haixiang and
               Mehta, Guneet S and Claus, Robert and Patel, Akhil J and Conklin,
               Matthew W and Inman, David R and Provenzano, Paolo P and Sifakis,
               Eftychios and Patel, Jignesh M and Eliceiri, Kevin W",
  journal   = "Front. Bioeng. Biotechnol.",
  publisher = "Frontiers Media SA",
  volume    =  8,
  pages     =  198,
  abstract  = "Quantification of fibrillar collagen organization has given new
               insight into the possible role of collagen topology in many
               diseases and has also identified candidate image-based
               bio-markers in breast cancer and pancreatic cancer. We have been
               developing collagen quantification tools based on the curvelet
               transform (CT) algorithm and have demonstrated this to be a
               powerful multiscale image representation method due to its unique
               features in collagen image denoising and fiber edge enhancement.
               In this paper, we present our CT-based collagen quantification
               software platform with a focus on new features and also giving a
               detailed description of curvelet-based fiber representation.
               These new features include C++-based code optimization for fast
               individual fiber tracking, Java-based synthetic fiber generator
               module for method validation, automatic tumor boundary generation
               for fiber relative quantification, parallel computing for
               large-scale batch mode processing, region-of-interest analysis
               for user-specified quantification, and pre- and post-processing
               modules for individual fiber visualization. We present a
               validation of the tracking of individual fibers and fiber
               orientations by using synthesized fibers generated by the
               synthetic fiber generator. In addition, we provide a comparison
               of the fiber orientation calculation on pancreatic tissue images
               between our tool and three other quantitative approaches. Lastly,
               we demonstrate the use of our software tool for the automatic
               tumor boundary creation and the relative alignment quantification
               of collagen fibers in human breast cancer pathology images, as
               well as the alignment quantification of in vivo mouse xenograft
               breast cancer images.",
  month     =  apr,
  year      =  2020,
  keywords  = "breast cancer; collagen organization; curvelet transform;
               fibrillar collagen; image analysis software; pancreatic cancer;
               second harmonic generation microscopy; tumor microenvironment",
  language  = "en"
}

@ARTICLE{Schlegel2023-gy,
  title    = "Tissue-engineered vocal fold replacement in swine: Methods for
              functional and structural analysis",
  author   = "Schlegel, Patrick and Yan, Kenneth and Upadhyaya, Sreenivasa and
              Buyens, Wim and Wong, Kirsten and Chen, Anthony and Faull, Kym F
              and Al-Hiyari, Yazeed and Long, Jennifer",
  journal  = "PLoS One",
  volume   =  18,
  number   =  4,
  pages    = "e0284135",
  abstract = "We have developed a cell-based outer vocal fold replacement (COVR)
              as a potential therapy to improve voice quality after vocal fold
              (VF) injury, radiation, or tumor resection. The COVR consists of
              multipotent human adipose-derived stem cells (hASC) embedded
              within a three-dimensional fibrin scaffold that resembles vocal
              fold epithelium and lamina propria layers. Previous work has shown
              improved wound healing in rabbit studies. In this pilot study in
              pigs, we sought to develop methods for large animal implantation
              and phonatory assessment. Feasibility, safety, and structural and
              functional outcomes of the COVR implant are described. Of eight
              pigs studied, six animals underwent COVR implantation with harvest
              between 2 weeks and 6 months. Recovery of laryngeal tissue
              structure was assessed by vibratory and histologic analyses.
              Recovery of voice function was assessed by investigating acoustic
              parameters that were derived specifically for pigs. Results showed
              improved lamina propria qualities relative to an injured control
              animal at 6 months. Acoustic parameters reflected voice worsening
              immediately after surgery as expected; acoustics displayed clear
              voice recovery in the animal followed for 6 months after COVR.
              These methods form the basis for a larger-scale long-term
              pre-clinical safety and efficacy study.",
  month    =  apr,
  year     =  2023,
  language = "en"
}

@ARTICLE{Shi2024-az,
  title     = "Novel bisphosphonate-based cathepsin {K}-triggered compound
               targets the enthesis without impairing soft tissue-to-bone
               healing",
  author    = "Shi, Brendan Y and Sriram, Varun and Wu, Shannon Y and Huang,
               Dave and Cheney, Alexis and Metzger, Melodie F and Sundberg,
               Oskar and Lyons, Karen M and McKenna, Charles E and Nishimura,
               Ichiro and Kremen, Jr, Thomas J",
  journal   = "Front. Bioeng. Biotechnol.",
  publisher = "Frontiers Media SA",
  volume    =  12,
  pages     =  1308161,
  abstract  = "Background: Osteoadsorptive fluorogenic sentinel 3 (OFS-3) is a
               recently described compound that contains a bone-targeting
               bisphosphonate (BP) and cathepsin K (Ctsk)-triggered fluorescence
               signal. A prior study in a murine Achilles repair model
               demonstrated its effectiveness at targeting the site of
               tendon-to-bone repair, but the intrinsic effect of this novel
               bisphosphonate chaperone on tendon-to-bone healing has not been
               previously explored. We hypothesized that application of this
               bisphosphonate-fluorophore cargo conjugate would not affect the
               biomechanical properties or histologic appearance of tendon-bone
               repairs. Materials and Methods: Right hindlimb Achilles
               tendon-to-bone repair was performed on 12-week old male mice.
               Animals were divided into 2 groups of 18 each: 1) Achilles repair
               with OFS-3 applied directly to the repair site prior to closure,
               and 2) Achilles repair with saline applied prior to closure.
               Repaired hindlimbs from 12 animals per group were harvested at 6
               weeks for biomechanical analysis with a custom 3D-printed jig. At
               4 and 6 weeks, repaired hindlimbs from the remaining animals were
               assessed histologically using H\&E, immunohistochemistry (IHC)
               staining for the presence of Ctsk, and second harmonic generation
               (SHG) imaging to evaluate collagen fibers. Results: At 6 weeks,
               there was no significant difference in failure load, stiffness,
               toughness, or displacement to failure between repaired hindlimbs
               that received OFS-3 versus saline. There was no difference in
               tissue healing on H\&E or Ctsk staining on immunohistochemistry
               between animals that received OFS-3 versus saline. Finally,
               second harmonic generation imaging demonstrated no difference in
               collagen fiber parameters between the two groups. Conclusion:
               OFS-3 did not significantly affect the biomechanical properties
               or histologic appearance of murine Achilles tendon-to-bone
               repairs. This study demonstrates that OFS-3 can target the site
               of tendon-to-bone repair without causing intrinsic negative
               effects on healing. Further development of this drug delivery
               platform to target growth factors to the site of tendon-bone
               repair is warranted.",
  month     =  feb,
  year      =  2024,
  keywords  = "biomechanics; enthesis; growth factor delivery; rotator cuff
               repair; targeted delivery",
  language  = "en"
}

@ARTICLE{Markus2023-rc,
  title     = "Effect of tissue fixation on the optical properties of structural
               components assessed by non-linear microscopy imaging",
  author    = "Markus, M Andrea and Ferrari, Daniele P and Alves, Frauke and
               Ramos-Gomes, Fernanda",
  journal   = "Biomed. Opt. Express",
  publisher = "Optica Publishing Group",
  volume    =  14,
  number    =  8,
  pages     = "3988--4002",
  abstract  = "Fixation methods such as formalin are commonly used for the
               preservation of tissue with the aim of keeping their structure as
               close as possible to the native condition. However, fixatives
               chemically interact with tissue molecules, such as collagen in
               the extracellular matrix (ECM) or myosin, and may thus modify
               their structure. Taking advantage of the second- and
               third-harmonic generation (SHG and THG) emission capabilities of
               such components, we used nonlinear two-photon microscopy (NL2PM)
               to evaluate the effect that preservation methods, such as
               chemical fixatives, have on the nonlinear capabilities of protein
               components within mouse tissues. Our results show that depending
               on the preservation technique used, the nonlinear capabilities of
               collagen, lipid droplets and myosin microarchitecture are
               strongly affected. Parameters of collagen fibers, such as density
               and branch points, especially in collagen-sparse regions, e.g.,
               in kidneys, were found to be altered upon formalin fixation.
               Moreover, cryo-freezing drastically reduced SHG signals from
               myosin. Our findings provide valuable information to select the
               best tissue fixation method for visualization and quantification
               of structural proteins, such as collagen and myosin by advanced
               NL2PM imaging techniques. This may advance the interpretation of
               the role these proteins play in disease.",
  month     =  aug,
  year      =  2023,
  keywords  = "Absorption coefficient;Imaging techniques;Nonlinear
               microscopy;Optical properties;Scattering media;Tissue optics",
  language  = "en"
}

@ARTICLE{Pence2021-zo,
  title     = "Multi-window sparse spectral sampling stimulated Raman scattering
               microscopy",
  author    = "Pence, Isaac J and Kuzma, Benjamin A and Brinkmann, Maximilian
               and Hellwig, Tim and Evans, Conor L",
  journal   = "Biomed. Opt. Express",
  publisher = "Optica Publishing Group",
  volume    =  12,
  number    =  10,
  pages     = "6095--6114",
  abstract  = "Stimulated Raman scattering (SRS) is a nondestructive and rapid
               technique for imaging of biological and clinical specimens with
               label-free chemical specificity. SRS spectral imaging is
               typically carried out either via broadband methods, or by tuning
               narrowband ultrafast light sources over narrow spectral ranges
               thus specifically targeting vibrational frequencies. We
               demonstrate a multi-window sparse spectral sampling SRS (S4RS)
               approach where a rapidly-tunable dual-output all-fiber optical
               parametric oscillator is tuned into specific vibrational modes
               across more than 1400 cm-1 during imaging. This approach is
               capable of collecting SRS hyperspectral images either by scanning
               a full spectrum or by rapidly tuning into select target
               frequencies, hands-free and automatically, across the
               fingerprint, silent, and high wavenumber windows of the Raman
               spectrum. We further apply computational techniques for spectral
               decomposition and feature selection to identify a sparse subset
               of Raman frequencies capable of sample discrimination. Here we
               have applied this novel method to monitor spatiotemporal dynamic
               changes of active pharmaceutical ingredients in skin, which has
               particular relevance to topical drug product delivery.",
  month     =  oct,
  year      =  2021,
  keywords  = "Biomedical imaging;Imaging systems;Imaging techniques;Optical
               parametric oscillators;Raman microscopy;Raman scattering",
  language  = "en"
}

@ARTICLE{Moffatt2026-qh,
  title     = "Radiation fibrosis of the epiglottis: Histopathology and
               multiphoton imaging of collagen and elastin",
  author    = "Moffatt, Clare and Pillutla, Pranati and Laklouk, Israa A and
               Smith, Alden F and Chhetri, Dinesh K",
  journal   = "Laryngoscope",
  publisher = "Wiley",
  volume    =  136,
  number    =  1,
  pages     = "340--350",
  abstract  = "OBJECTIVES: Dysphagia attributed to radiation-induced fibrosis
               (RIF) is common among patients treated with radiation for head
               and neck cancer. In this study, we characterize RIF in the
               epiglottis. This has not been previously described. METHODS: Our
               cohort included 26 patients who underwent partial
               epiglottidectomy for symptomatic dysphagia due to epiglottic
               dysfunction. Eighteen patients had a history of radiation
               therapy. Epiglottis tissue sections were stained with H\&E, and
               the ratio of epiglottic cartilage thickness to total epiglottis
               thickness was measured. Imaging for elastin and collagen was
               performed with multiphoton microscopy, and the density of each in
               the lamina propria layer was quantified. RESULTS: Radiated
               epiglottis had a higher prevalence of dilated vasculature,
               chronic inflammation, focal hyalinization-like changes in the
               elastic cartilage, and atrophic seromucous glands. In addition, a
               lower ratio of elastic cartilage to total epiglottis thickness
               was found (0.36 in radiated versus 0.45 in non-radiated, p <
               0.01), attributable to increased thickness of the lamina propria
               layer due to fibrosis. In the radiated samples, the density of
               collagen was higher, 46.51\% versus 36.78\% (p < 0.01), and the
               density of elastin was lower, 5.11\% versus 8.27\% (p < 0.01).
               Elastic fibers were straighter, shorter, and more fragmented in
               radiated samples. CONCLUSION: This first characterization of RIF
               and pilot use of multiphoton microscopy in the epiglottis
               identified key histopathologic characteristics associated with
               radiation, notably higher density of collagen, lower density of
               elastin, and fragmentation of elastic fibers. These findings
               provide a pathophysiologic explanation for the relationship
               between epiglottic dysfunction, dysphagia, and radiation. LEVEL
               OF EVIDENCE: Level 3.",
  month     =  jan,
  year      =  2026,
  keywords  = "collagen; elastin; epiglottis; radiation fibrosis",
  language  = "en"
}

@ARTICLE{Li2025-wb,
  title    = "Arterial compliance is improved via enteral serine protease
              inhibition in experimental trauma/hemorrhagic shock",
  author   = "Li, Joyce B and Dos Santos, Fernando and Muller, Cynthia R and
              Moreira, Nathalia J D and Borges, Luciano F and Irigoyen, Maria
              Claudia C and Schmid-Schönbein, Geert W and Kistler, Erik B",
  journal  = "Front. Physiol.",
  volume   =  16,
  pages    =  1655946,
  abstract = "Background: Systemic hypotension remains a challenge in
              trauma/hemorrhagic shock (T/HS). Despite intensive vasopressor and
              fluid therapy, mean arterial blood pressure (MAP) may become
              refractory to treatment. Arterial compliance (AC) is a critical
              determinant of arterial hemodynamics but is often overlooked in
              acute shock states. Considering previous findings on the benefits
              of enteral protease inhibition in preserving vascular resistance
              after T/HS, this study investigated both the role of AC and the
              effects of enteral protease inhibition on AC in T/HS. Methods:
              Wistar rats underwent experimental T/HS by laparotomy and
              exsanguination to induce a MAP of ∼40 mmHg for 90 min. Animals
              were randomized into three groups corresponding to the
              intervention: shed whole blood (WB), Lactated Ringer's (LR), and
              LR with enteral gabexate mesilate treatment (LR+GM). Resuscitation
              (120-min period) was initiated by fluid reperfusion with a goal
              MAP of 65 mmHg. AC was measured via pulse wave velocity (PWV),
              passive pressure myography, and atomic force microscopy (AFM),
              with healthy donor arteries for comparison. Results: PWV increased
              by ∼15\% in all groups after shock. After resuscitation, LR-only
              animals maintained high PWVs, but significantly lower diastolic
              pressures (27 mmHg) compared to GM-treated (37 mmHg; p < 0.05) and
              those reperfused with WB (52 mmHg; p < 0.01). T/HS arteries,
              particularly the untreated LR arteries, exhibited leftward shifts
              in circumferential tension-strain curves. LR arteries exhibited
              higher tangent moduli (5 N/m; p < 0.01) at low physiological
              stresses, which was corroborated by reduced opening angles,
              increased mechanical stiffness, alterations in the extracellular
              matrix, and increased MMP/elastase-like activity. LR+GM and WB
              arteries displayed elastic moduli and vascular structures more
              similar to those of healthy arteries. Conclusion: Experimental
              T/HS results in impaired AC, which is partially attenuated by
              enteral GM administration. Vascular biomechanical impairment may
              underlie the unrestored MAP in fulminant shock. By targeting
              modulators of AC, with enteral serine protease inhibition as an
              adjunct intervention, hemodynamic stability and patient outcomes
              may be improved in T/HS.",
  month    =  aug,
  year     =  2025,
  keywords = "arterial compliance; circulatory shock; diastolic pressure;
              hemorrhagic shock; protease inhibition; resuscitation; trauma",
  language = "en"
}

@ARTICLE{Prevedel2019-ve,
  title     = "Brillouin microscopy: an emerging tool for mechanobiology",
  author    = "Prevedel, Robert and Diz-Muñoz, Alba and Ruocco, Giancarlo and
               Antonacci, Giuseppe",
  journal   = "Nat. Methods",
  publisher = "Springer Science and Business Media LLC",
  volume    =  16,
  number    =  10,
  pages     = "969--977",
  abstract  = "The role and importance of mechanical properties of cells and
               tissues in cellular function, development and disease has widely
               been acknowledged, however standard techniques currently used to
               assess them exhibit intrinsic limitations. Recently, Brillouin
               microscopy, a type of optical elastography, has emerged as a
               non-destructive, label- and contact-free method that can probe
               the viscoelastic properties of biological samples with
               diffraction-limited resolution in 3D. This led to increased
               attention amongst the biological and medical research
               communities, but it also sparked debates about the interpretation
               and relevance of the measured physical quantities. Here, we
               review this emerging technology by describing the underlying
               biophysical principles and discussing the interpretation of
               Brillouin spectra arising from heterogeneous biological matter.
               We further elaborate on the technique's limitations, as well as
               its potential for gaining insights in biology, in order to guide
               interested researchers from various fields.",
  month     =  oct,
  year      =  2019,
  language  = "en"
}

@ARTICLE{Cisek2021-fy,
  title     = "Polarization-sensitive second harmonic generation microscopy for
               investigations of diseased collagenous tissues",
  author    = "Cisek, Richard and Joseph, Ariana and Harvey, Macaulay and
               Tokarz, Danielle",
  journal   = "Front. Phys.",
  publisher = "Frontiers Media SA",
  volume    =  9,
  abstract  = "The advancement of non-invasive quantitative optical diagnosis
               techniques such as polarization-sensitive second harmonic
               generation microscopy (PSHG) for diseases such as cancer presents
               opportunities for improving disease understanding and survival
               rates. Here, novel and developing techniques in PSHG microscopy
               applied for the differentiation of cancerous or diseased tissues
               are presented, including circular dichroism, modulation of laser
               linear polarization, detection of outgoing linear laser
               polarization, and double-Stokes Mueller. Typically, initial
               cancer diagnosis is performed by visual inspection of stained
               biopsy or surgical resection tissue sections under bright-field
               microscopy, however, early diagnosis is challenging due to
               variability in morphological interpretation of the tissues, and
               because cancer initiation regions can be small and easy to miss.
               Therefore, pathologists could benefit in identifying cancer on
               biopsy or surgical resection sections by using unbiased
               quantitative automated technologies with high spatial resolution
               and improved disease specificity that can check the entire slide
               pixel-by-pixel. Second harmonic generation microscopy offers the
               opportunity to measure ultrastructural alterations in collagenous
               scaffolds of organ tissues virtually background free on
               submicron-sized tissue regions. The approach is particularly
               interesting for cancer diagnosis applications, because during
               cancer initiation and progression, the collagen in the affected
               tissue extracellular matrix is often deregulated and becomes
               disorganized. This mini-review contains a thorough summary of
               PSHG techniques that have interrogated diseased tissues, and
               discusses their technical variations and successes in disease
               discrimination.",
  month     =  aug,
  year      =  2021
}

@ARTICLE{Burkitt2025-se,
  title     = "Multiphoton luminescence imaging: A label-free tool for
               visualizing the long-term fate of gold nanoparticles in tissue",
  author    = "Burkitt, Sean and Eremina, Olga E and Mouchawar, Anthony and
               Fernando, Augusta and Edmondson, Elijah and Stern, Stephan and
               Marks, Carolyn and Zavaleta, Cristina",
  journal   = "ACS Nano",
  publisher = "American Chemical Society (ACS)",
  volume    =  19,
  number    =  32,
  pages     = "29286--29300",
  abstract  = "Gold nanoparticles (AuNPs) are increasingly used in applications
               across the biomedical domain, yet their long-term biodistribution
               and biocompatibility remain poorly understood. Conventional
               brightfield microscopy imaging techniques often fail to detect
               AuNPs due to optical diffraction limits and lack of chromogenic
               contrast. Understanding the biodistribution and ultimate fate of
               these nonbiodegradable NPs is crucial for further development of
               AuNP-based therapeutics and diagnostics. Here, we present a
               label-free multiphoton luminescence (MPL) imaging workflow that
               enables sensitive detection of AuNPs in liver histology sections,
               even 1 year after intravenous (IV) administration. MPL imaging
               exploits the intrinsic nonlinear optical properties of AuNPs to
               generate broadband emission under ultrafast pulsed laser
               excitation, enabling subcellular localization without exogenous
               labels while having the ability to rapidly image entire organ
               sections. The intrinsic, distinct broadband MPL emission produced
               by gold allows us to study these NPs in their biological context
               without extrinsic labels while also faithfully representing the
               surrounding tissue architecture via autofluorescence and second
               harmonic generation. We demonstrate that MPL imaging detects up
               to 98\% more AuNP-positive regions than brightfield microscopy in
               challenging low-dose (1 nM) conditions and requires no
               modification of standard histology workflows. Correlative imaging
               with SEM-EDS confirms high spatial specificity (AUC = 0.955) of
               MPL for AuNP localization. Dose-dependent retention patterns were
               observed across liver tissue, and MPL analysis showed strong
               correlation with ICP-MS quantification. Importantly, histological
               and immunohistochemical analyses (Masson's trichrome, CD3, and
               TUNEL) revealed no significant fibrosis, immune activation, or
               apoptosis in liver tissue at either low (1 nM) or high (10 nM)
               doses at 1 year post IV administration. These findings establish
               MPL imaging as a robust, label-free tool for long-term tracking
               of AuNPs in biological tissue and highlight its potential for
               improving biodistribution and safety assessments.",
  month     =  aug,
  year      =  2025,
  keywords  = "biodistribution; digital pathology; gold nanoparticles;
               multiphoton imaging; multiphoton luminescence",
  language  = "en"
}

@ARTICLE{Kashani-Ligumsky2026-nt,
  title     = "Placenta accreta spectrum: disrupted collagen architecture at a
               previous scar is a defining characteristic of placental adherence",
  author    = "Kashani Ligumsky, Lior and Jeong, Anhyo and Martinez, Guadalupe
               and Marks, Haley and Shah, Sohum and Staniczek, Jakub and Smith,
               Caroline E and Shainker, Scott A and Karumanchi, S Ananth and
               Bentolila, Laurent A and Krakow, Deborah and Megli, Christina J
               and Afshar, Yalda",
  journal   = "Am. J. Obstet. Gynecol.",
  publisher = "Elsevier BV",
  volume    =  233,
  number    = "6S",
  pages     = "S645--S661.e1",
  abstract  = "BACKGROUND: Placenta accreta spectrum is a severe obstetric
               complication associated with uterine scarring, particularly from
               prior cesarean births. Despite its considerable clinical
               sequalae, the underlying mechanisms remain unclear. Recent
               evidence suggests that abnormal extracellular matrix remodeling
               plays a pivotal role in placenta accreta spectrum pathology.
               OBJECTIVE: This study investigates collagen architecture and its
               role in placenta accreta spectrum adherence using multimodal
               imaging, an in vivo mouse model, and an in vitro co-culture
               system, aiming to identify pathophysiology of placenta accreta
               spectrum with implications for improved uterine repair and
               placenta accreta spectrum prevention. STUDY DESIGN: Human
               placenta accreta spectrum and nonplacenta accreta spectrum
               specimens were analyzed using nonlinear label-free optical
               imaging techniques to characterize collagen architecture at the
               decidual-placental interface (total n=23, placenta accreta
               spectrum=13). A surgical mouse model of placenta accreta spectrum
               was used to examine collagen organization and placental adherence
               patterns. Additionally, an in vitro ``accreta-in-a-dish'' model
               was developed using decidualized human uterine fibroblasts,
               trophoblasts, and macrophages to assess extracellular matrix
               remodeling, wound healing, and inflammatory interactions.
               Live-cell imaging and electric cell-substrate impedance sensing
               quantified the impact of collagen deposition and inflammation on
               scar resistance to electric impedance and trophoblast behavior.
               RESULTS: Disorganized fibrillar collagen deposition with
               disrupted border integrity was a hallmark of placenta accreta
               spectrum pathology, specifically at the site of adherent,
               compared to nonadherent placenta accreta spectrum controls in
               human surgical specimens. The mouse model of placenta accreta
               spectrum confirmed matrix disorganization at the site of abnormal
               placental adherence. In vitro, collagen coating enhanced wound
               healing, while placental-conditioned media from placenta accreta
               spectrum sites impaired macrophage-driven tissue repair. In
               vitro, trophoblasts preferentially avoided extracellular
               matrix-rich regions, suggesting a role for matrix composition in
               placental attachment and invasion. Macrophage-mediated
               inflammation further compromised scar resistance, indicating an
               inflammatory-matrix interplay that may predispose to placenta
               accreta spectrum. CONCLUSION: These findings underscore the role
               of extracellular matrix dysregulation in placenta accreta
               spectrum, highlighting collagen architecture as a critical
               determinant of placental adherence. Collagen modulation and
               targeted inflammatory interventions could improve uterine scar
               healing and reduce placenta accreta spectrum incidence. Future
               research should focus on translating these mechanistic insights
               into diagnostic and therapeutic strategies for placenta accreta
               spectrum prevention and management.",
  month     =  jan,
  year      =  2026,
  language  = "en"
}

@ARTICLE{Kremen2023-et,
  title     = "Biologically-coupled bisphosphonate chaperones effectively
               deliver molecules to the site of soft tissue-bone healing",
  author    = "Kremen, Jr, Thomas J and Shi, Brendan Y and Wu, Shannon Y and
               Sundberg, Oskar and Sriram, Varun and Kim, Won and Sheyn, Dmitriy
               and Lyons, Karen M and Wang, Weiguang and McKenna, Charles E and
               Nishimura, Ichiro",
  journal   = "J. Orthop. Res.",
  publisher = "John Wiley \& Sons, Ltd",
  volume    =  41,
  number    =  10,
  pages     = "2250--2260",
  abstract  = "Tendon injuries are common and often treated surgically, however,
               current tendon repair healing results in poorly organized
               fibrotic tissue. While certain growth factors have been reported
               to improve both the strength and organization of the repaired
               enthesis, their clinical applicability is severely limited due to
               a lack of appropriate delivery strategies. In this study, we
               evaluated a recently developed fluorescent probe, Osteoadsorptive
               Fluorogenic Sentinel-3 that is composed of a bone-targeting
               bisphosphonate (BP) moiety linked to fluorochrome and quencher
               molecules joined via a cathepsin K-sensitive peptide sequence.
               Using a murine Achilles tendon-to-bone repair model, BP-based
               and/or Ctsk-coupled imaging probes were applied either locally or
               systemically. Fluorescence imaging was used to quantify the
               resultant signal in vivo. After tendon-bone repair, animals that
               received either local or systemic administration of imaging
               probes demonstrated significantly higher fluorescence signal at
               the repair site compared to the sham surgery group at all time
               points (p < 0.001), with signal peaking at 7-10 days after
               surgery. Our findings demonstrate the feasibility of using a
               novel BP-based targeting and Ctsk-activated delivery of molecules
               to the site of tendon-to-bone repair and creates a foundation for
               further development of this platform as an effective strategy to
               deliver bioactive molecules to sites of musculoskeletal injury.",
  month     =  oct,
  year      =  2023,
  keywords  = "cell and molecular imaging; surgical repair; tendon",
  language  = "en"
}

@ARTICLE{Arshee2025-wg,
  title     = "Impact of paraben on uterine collagen: An integrated and targeted
               Correlative approach using second harmonic generation microscopy,
               nanoindentation, and atomic force microscopy",
  author    = "Arshee, Mahmuda R and Shukla, Ritwik and Li, Jie and Doha, Umnia
               and Bagchi, Indrani C and Ziv-Gal, Ayelet and Wagoner Johnson,
               Amy J",
  journal   = "J. Mech. Behav. Biomed. Mater.",
  publisher = "Elsevier BV",
  volume    =  165,
  number    =  106926,
  pages     =  106926,
  abstract  = "This study investigates the structural and mechanical changes in
               uterine collagen following exposure to propylparaben (PP) using a
               combined methodology of Second Harmonic Generation (SHG)
               microscopy, Nanoindentation (NI), and Atomic Force Microscopy
               (AFM). SHG analysis identified significant disorganization in
               collagen fibril orientation in the circumferential layer and
               heterogeneous distribution of regions with elevated forward to
               backward ratios (F/B) across all uterine layers due to PP
               exposure. High F/B can indicate multiple potential fibril-level
               changes like thickened fibrils, higher crosslinking, fibril
               disorganization - changes not fully decipherable by SHG alone.
               Recognizing this limitation, the study employs NI and AFM to
               provide complementary mechanical and nanoscale insights. NI
               revealed increased indentation modulus in the exposed uteri,
               suggesting increased stiffness. Co-registration of the
               indentation response with SHG parameters uncovered that elevated
               F/B regions show enhanced mechanical stiffness, suggesting a
               fibrotic transformation following chronic PP exposure. AFM was
               specifically performed on regions identified by SHG as having low
               or high F/B, providing the necessary nanoscale resolution to
               elucidate the structural changes in fibrils that are likely
               responsible for the observed alterations. AFM confirmed the
               presence of disordered and entangled collagen fibrils in the
               circumferential layer in all regions and an increase in fibril
               diameter in the high F/B regions in the PP-exposed uteri.
               Together, these findings demonstrate significant alterations in
               collagen architecture due to PP exposure, revealing disruptions
               at both the fiber and fibril levels and highlighting the
               potential for broader applications of the multi-scale,
               multi-modal approach in collagenous tissue studies.",
  month     =  may,
  year      =  2025,
  keywords  = "Atomic force microscopy; Collagen; Indentation; Paraben; Second
               harmonic generation (SHG); Uterus",
  language  = "en"
}

@ARTICLE{de-Lizaraga2025-gc,
  title     = "Probing the role of telopeptides in collagen fibrillogenesis by
               second harmonic scattering",
  author    = "de Lizaraga, Joshua and Martin, Marta and Salmon, Estelle and
               Cloitre, Thierry and Gergely, Csilla and Jonin, Christian",
  journal   = "J. Phys. Chem. B",
  publisher = "American Chemical Society (ACS)",
  volume    =  129,
  number    =  35,
  pages     = "9020--9029",
  abstract  = "Collagen, the primary structural protein in the extracellular
               matrix, plays a critical role in tissue architecture and
               mechanical integrity. This study investigates the structural and
               nonlinear optical properties of atelocollagen (Acol) and
               telocollagen (Tcol) in response to pH variations by using second
               harmonic scattering (SHS), polarization-resolved SHS (P-SHS), and
               atomic force microscopy (AFM). AFM imaging revealed distinct
               morphological differences, with Acol forming wavy,
               non-cross-linked, and randomly arranged fibers, while Tcol
               exhibited a more interconnected, mesh-like fibrillar network. SHS
               and P-SHS measurements further confirmed these differences,
               indicating significant variations in hyperpolarizability and
               depolarization ratio (DR) between Acol and Tcol. The results
               suggest that telopeptides play a crucial role in collagen
               fibrillogenesis, influencing the molecular symmetry and optical
               responses. The observed differences in nonlinear optical
               properties highlight the potential of SHS as a powerful tool for
               characterizing collagen self-assembly mechanisms. These findings
               contribute to a broader understanding of collagen organization in
               biological and biomaterial contexts, with implications for tissue
               engineering, fibrosis research, and regenerative medicine. Future
               work should explore how external factors, such as ionic strength
               and cross-linking agents, further modulate collagen fibril
               formation and its mechanical properties.",
  month     =  sep,
  year      =  2025,
  language  = "en"
}

@ARTICLE{Khattignavong2023-xs,
  title     = "Development of a facile method to compute collagen network
               pathological anisotropy using {AFM} imaging",
  author    = "Khattignavong, Emilie and Neshatian, Mehrnoosh and Vaez, Mina and
               Guillermin, Amaury and Tauer, Josephine T and Odlyha, Marianne
               and Mittal, Nimish and Komarova, Svetlana V and Zahouani, Hassan
               and Bozec, Laurent",
  journal   = "Sci. Rep.",
  publisher = "Springer Science and Business Media LLC",
  volume    =  13,
  number    =  1,
  pages     =  20173,
  abstract  = "Type I collagen, a fundamental extracellular matrix (ECM)
               component, is pivotal in maintaining tissue integrity and
               strength. It is also the most prevalent fibrous biopolymer within
               the ECM, ubiquitous in mammalian organisms. This structural
               protein provides essential mechanical stability and resilience to
               various tissues, including tendons, ligaments, skin, bone, and
               dentin. Collagen has been structurally investigated for several
               decades, and variation to its ultrastructure by histology has
               been associated with several pathological conditions. The current
               study addresses a critical challenge in the field of collagen
               research by providing a novel method for studying collagen fibril
               morphology at the nanoscale. It offers a computational approach
               to quantifying collagen properties, enabling a deeper
               understanding of how collagen type I can be affected by
               pathological conditions. The application of Fast Fourier
               Transform (FFT) coupled with Atomic Force Microscope (AFM)
               imaging distinguishes not only healthy and diseased skin but also
               holds potential for automated diagnosis of connective tissue
               disorders (CTDs), contributing to both clinical diagnostics and
               fundamental research in this area. Here we studied the changes in
               the structural parameters of collagen fibrils in Ehlers Danlos
               Syndrome (EDS). We have used skin extracted from genetically
               mutant mice that exhibit EDS phenotype as our model system
               (Col1a1Jrt/+ mice). The collagen fibrils were analyzed by AFM
               based descriptive-structural parameters, coupled with a 2D Fast
               Fourier Transform(2D-FFT) approach that automated the analysis of
               AFM images. In addition, each sample was characterized based on
               its FFT and power spectral density. Our qualitative data showed
               morphological differences in collagen fibril clarity (clearness
               of the collagen fibril edge with their neighbouring fibri),
               D-banding, orientation, and linearity. We have also demonstrated
               that FFT could be a new tool for distinguishing healthy from
               tissues with CTDs by measuring the disorganization of fibrils in
               the matrix. We have also employed FFT to reveal the orientations
               of the collagen fibrils, providing clinically relevant phenotypic
               information on their organization and anisotropy. The result of
               this study can be used to develop a new automated tool for better
               diagnosis of CTDs.",
  month     =  nov,
  year      =  2023,
  language  = "en"
}

@ARTICLE{Jones2022-zk,
  title     = "Structurally abnormal collagen fibrils in abdominal aortic
               aneurysm resist platelet adhesion",
  author    = "Jones, Blain and Debski, Anna and Hans, Chetan P and Go, Michael
               R and Agarwal, Gunjan",
  journal   = "J. Thromb. Haemost.",
  publisher = "Elsevier BV",
  volume    =  20,
  number    =  2,
  pages     = "470--477",
  abstract  = "BACKGROUND: Platelet adhesion to the subendothelial collagen
               fibrils is one of the first steps in hemostasis. Understanding
               how structural perturbations in the collagen fibril affect
               platelet adhesion can provide novel insights into disruption of
               hemostasis in various diseases. We have recently identified the
               presence of abnormal collagen fibrils with compromised D-periodic
               banding in the extracellular matrix remodeling present in
               abdominal aortic aneurysms (AAA). OBJECTIVE: In this study, we
               employed multimodal microscopy approaches to characterize how
               collagen fibril structure impacts platelet adhesion in clinical
               AAA tissues. METHODS: Ultrastructural atomic force microscopy
               (AFM) analysis was performed on tissue sections after staining
               with fluorescently labeled collagen hybridizing peptide (CHP) to
               recognize degraded collagen. Second harmonic generation (SHG)
               microscopy was used on CHP-stained sections to identify regions
               of intact versus degraded collagen. Finally, platelet adhesion
               was identified via SHG and indirect immunofluorescence on the
               same tissue sections. RESULTS: Our results indicate that
               ultrastructural features characterizing collagen fibril
               abnormalities coincide with CHP staining. SHG signal was absent
               from CHP-positive regions. Additionally, platelet binding was
               primarily localized to regions with SHG signal. Abnormal collagen
               fibrils present in AAA (in SHG negative regions) were thus found
               to inhibit platelet adhesion compared to normal fibrils.
               CONCLUSIONS: Our investigations reveal how the collagen fibril
               structure in the vessel wall can serve as another regulator of
               platelet-collagen adhesion. These results can be broadly applied
               to understand the role of collagen fibril structure in regulating
               thrombosis or bleeding disorders.",
  month     =  feb,
  year      =  2022,
  keywords  = "abdominal aortic aneurysm; atomic force microscopy; blood
               platelets; collagen; fluorescence; microscopy",
  language  = "en"
}

@ARTICLE{MacArthur2025-vv,
  title     = "Quantifying collagen ultrastructural changes in human lumbar disc
               degeneration using polarization resolved second harmonic
               generation microscopy",
  author    = "MacArthur, Sasha Dawn",
  publisher = "Halifax, NS: Saint Mary's University",
  abstract  = "Intervertebral disc degeneration (IVD) involves ultrastructural
               disruption of the collagen network, yet conventional imaging
               cannot resolve these early changes. This thesis applies
               polarization-resolved second harmonic generation (PSHG)
               microscopy to quantify collagen organization in human lumbar IVD
               tissue across degeneration grades. A preliminary porcine study
               demonstrated that PSHG provides stable, reproducible parameter
               estimation in the annulus fibrosus (AF), motivating its
               application to human samples. Using PSHG, multiple parameters
               were evaluated, including collagen disorder (ρ), chirality (κ),
               and degree of linear polarization, along with associated
               distribution widths and variability. Several SHG-derived collagen
               parameters changed with degeneration. In discs without vertebral
               endplate changes (Modic 0), these parameters distinguished
               between moderate and advanced degeneration (Pfirrmann III vs IV).
               In contrast, in discs with Modic 2 changes, differences between
               Pfirrmann grades were reduced, suggesting that collagen
               structural alterations occur prior to vertebral endplate
               pathology and are already established once Modic changes are
               present.",
  year      =  2025
}

@ARTICLE{Plekhanov2023-yb,
  title     = "Side-by-side {OCE}-study of elasticity and {SHG}-characterization
               of collagen fibers in breast cancer tissue before and after
               chemotherapy",
  author    = "Plekhanov, Anton A and Potapov, Arseniy L and Pavlov, Mikhail V
               and Elagin, Vadim V and Gubarkova, Ekaterina V and Sovetsky,
               Alexander A and Matveev, Lev A and Vorontsov, Dmitriy A and
               Matveyev, Alexander L and Vorontsov, Alexey Y and Gamayunov,
               Sergey V and Zagaynova, Elena V and Sirotkina, Marina A and
               Zaitsev, Vladimir Y and Gladkova, Natalia D",
  journal   = "J. Biomed. Photonics Eng.",
  publisher = "Samara National Research University",
  address   = "Россия, Самара",
  volume    =  9,
  number    =  2,
  pages     =  020305,
  abstract  = "In the recent years promising results have been shown by the use
               of Compression optical coherence elastography (C-OCE) as a new
               optical biopsy approach to morphological assessment/diagnostics
               of human breast cancer using differences in elastic properties of
               morphological components of cancerous tissues. In this study, for
               the first time, a relationship was established between
               microstructural organization and biomechanical properties of
               breast-cancer tissue with pathomorphological changes caused by
               chemotherapy. To characterize texture of collagen fibers in the
               microenvironment of breast cancer, high-resolution visualization
               by Second-harmonic generation (SHG) microscopy was used. A
               side-by-side C-OCE and SHG imaging of patients’ breast cancer
               tissues before and after chemotherapy was carried out. Regions of
               the cancer stroma (collagen fibers outside aggregates of cancer
               cells) were assessed separately from regions of cancer cell
               clusters penetrated by collagen fibers. For cancer stroma areas
               after chemotherapy, a statistically significant decrease in
               stiffness values was found. Simultaneously, parameters of
               collagen texture in SHG images (mean intensity, “coherency” and
               “energy”) indicated increase in the collagen content,
               orientational orderliness, and collagen-texture heterogeneity. In
               contrast, cancer-cell areas post chemotherapy showed a
               statistically significant increase in stiffness. Analysis of SHG
               images of these regions indicated decrease in the inter-cellular
               collagen content and heterogeneity of its texture, whereas its
               orientational orderliness somewhat increased. The established
               negative correlation between stiffness and SHG parameters of
               collagen in cancer stroma indicates the contribution of the
               increase in orientational orderliness and total collagen content
               to the reduction in stiffness of breast cancer stroma after
               chemotherapy. For cancer-cell regions, significantly lower
               correlation between stiffness and SHG parameters (especially for
               coherency) was found, indicating stronger role of
               chemotherapy-induced changes in cancer cells themselves. These
               results give a deeper insight in the role of collagen texture
               organization in biomechanics of breast cancer tissues and
               contribute to a more detailed substantiation of the morphological
               characterization of breast cancer by C-OCE imaging. © 2023
               Journal of Biomedical Photonics \& Engineering.",
  month     =  jun,
  year      =  2023,
  keywords  = "breast cancer, tissue stiffness, morphological assessment,
               compression optical coherence elastography (C-OCE),
               second-harmonic generation (SHG); breast cancer, tissue
               stiffness, morphological assessment, compression optical
               coherence elastography (C-OCE), second-harmonic generation (SHG)"
}

@ARTICLE{Harvey2023-kw,
  title     = "Effect of out of plane orientation on polarization second
               harmonic generation of single collagen fibrils",
  author    = "Harvey, M A and Cisek, R and Tokarz, D and {others}",
  journal   = "Biomed. Opt. Express",
  publisher = "opg.optica.org",
  year      =  2023
}

@ARTICLE{Lopez-De-Padilla2021-hb,
  title     = "Picrosirius red staining: Revisiting its application to the
               qualitative and quantitative assessment of collagen type {I} and
               type {III} in tendon",
  author    = "López De Padilla, Consuelo M and Coenen, Michael J and Tovar,
               Alejandro and De la Vega, Rodolfo E and Evans, Christopher H and
               Müller, Sebastian A",
  journal   = "J. Histochem. Cytochem.",
  publisher = "SAGE Publications",
  volume    =  69,
  number    =  10,
  pages     = "633--643",
  abstract  = "Collagen has a major role in the structural organization of
               tendons. Picrosirius red (PSR) staining viewed under polarized
               light microscopy is the standard method to evaluate the
               organization of collagen fibers in tissues. It is also used to
               distinguish between type I and type III collagen in tissue
               sections. However, accurate analysis and interpretation of PSR
               images are challenging because of technical factors and
               historical misconceptions. The aim of this study was to clarify
               whether collagen types I and III can be distinguished by PSR
               staining in rat Achilles tendons, using double
               immunohistochemistry as the positive control. Our findings showed
               that PSR staining viewed with polarized light microscopy was
               suitable for qualitative and quantitative assessment of total
               collagen but was not able to distinguish collagen types. We found
               it critical to use a polarizing microscope equipped with a
               rotating stage; tendon section orientation at 45° with respect to
               crossed polarizers was optimal for the qualitative and
               quantitative assessment of collagen organization.
               Immunohistochemistry was superior to PSR staining for detection
               of collagen type III. We also compared formalin and Bouin
               solution as fixatives. Both produced similar birefringence, but
               formalin-fixed tendons provided higher quality histological
               detail with both hematoxylin-eosin and immunostaining.",
  month     =  oct,
  year      =  2021,
  keywords  = "Achilles tendon; Sirius red; collagen type; immunostaining;
               microscopy; polarization",
  language  = "en"
}

@ARTICLE{Cho2023-rt,
  title     = "Atomic force microscopy-mediated mechanobiological profiling of
               complex human tissues",
  author    = "Cho, David H and Aguayo, Sebastian and Cartagena-Rivera,
               Alexander X",
  journal   = "Biomaterials",
  publisher = "Elsevier BV",
  volume    =  303,
  number    =  122389,
  pages     =  122389,
  abstract  = "Tissue mechanobiology is an emerging field with the overarching
               goal of understanding the interplay between biophysical and
               biochemical responses affecting development, physiology, and
               disease. Changes in mechanical properties including stiffness and
               viscosity have been shown to describe how cells and tissues
               respond to mechanical cues and modify critical biological
               functions. To quantitatively characterize the mechanical
               properties of tissues at physiologically relevant conditions,
               atomic force microscopy (AFM) has emerged as a highly versatile
               biomechanical technology. In this review, we describe the
               fundamental principles of AFM, typical AFM modalities used for
               tissue mechanics, and commonly used elastic and viscoelastic
               contact mechanics models to characterize complex human tissues.
               Furthermore, we discuss the application of AFM-based
               mechanobiology to characterize the mechanical responses within
               complex human tissues to track their developmental,
               physiological/functional, and diseased states, including oral,
               hearing, and cancer-related tissues. Finally, we discuss the
               current outlook and challenges to further advance the field of
               tissue mechanobiology. Altogether, AFM-based tissue
               mechanobiology provides a mechanistic understanding of biological
               processes governing the unique functions of tissues.",
  month     =  dec,
  year      =  2023,
  language  = "en"
}

@ARTICLE{Binnig1986-xy,
  title     = "Atomic force microscope",
  author    = "Binnig, G and Quate, C F and Gerber, C",
  journal   = "Phys. Rev. Lett.",
  publisher = "American Physical Society (APS)",
  volume    =  56,
  number    =  9,
  pages     = "930--933",
  month     =  mar,
  year      =  1986,
  language  = "en"
}

@ARTICLE{Plodinec2010-wg,
  title     = "Imaging articular cartilage tissue using atomic force microscopy
               ({AFM})",
  author    = "Plodinec, Marija and Loparic, Marko and Aebi, Ueli",
  journal   = "Cold Spring Harb. Protoc.",
  publisher = "Cold Spring Harbor Laboratory",
  volume    =  2010,
  number    =  10,
  pages     = "db.prot5499",
  abstract  = "Cartilage is a complex avascular tissue composed of cells
               (``chondrocytes'') embedded in an extracellular matrix (ECM)
               consisting of 70\%-80\% water. The primary components of the ECM
               are negatively charged aggrecans and collagen II fibrils, which
               possess a characteristic, ordered three-dimensional structure.
               The components interact to ensure that the cartilage is able to
               absorb shock and can function to protect the bone ends. Atomic
               force microscopy (AFM) can be used to examine structure-function
               relationships of cartilage at both micrometer and nanometer
               scales. When imaged at the micrometer scale with microspheres,
               only the ECM and chondrocytes can be distinguished.
               Correspondingly, mechanical testing of cartilage at the
               micrometer scale results in unimodal distribution of the
               stiffness because the bulk elastic property of the ECM is probed.
               In contrast, bare AFM tips are able to reveal the molecular
               components of the ECM at the nanometer scale. Mechanical testing
               at the nanometer scale reveals a bimodal distribution of the
               stiffness and reflects the distinct stiffness of the collagen
               network and the proteoglycan moiety. In this protocol, the
               corresponding AFM image and force map are used to reveal the
               distinct morphology of the collagen fibers and proteoglycan gel.
               Although, in principle, these experiments can be performed using
               any AFM, an AFM with tube scanners that have manual screws for
               tilting the sample is preferable because cartilage has
               macroscopically rough surface features. By manually tilting the
               probe over the sample, an optimal angle for tip approach can be
               achieved.",
  month     =  oct,
  year      =  2010,
  language  = "en"
}

@ARTICLE{Quansah2023-um,
  title     = "Investigating biochemical and structural changes of glycated
               collagen using multimodal multiphoton imaging, Raman
               spectroscopy, and atomic force microscopy",
  author    = "Quansah, Elsie and Shaik, Tanveer Ahmed and Çevik, Ecehan and
               Wang, Xinyue and Höppener, Christiane and Meyer-Zedler, Tobias
               and Deckert, Volker and Schmitt, Michael and Popp, Jürgen and
               Krafft, Christoph",
  journal   = "Anal. Bioanal. Chem.",
  publisher = "Springer Science and Business Media LLC",
  volume    =  415,
  number    =  25,
  pages     = "6257--6267",
  abstract  = "Advanced glycation end products (AGEs) form extracellular
               crosslinking with collagenous proteins, which contributes to the
               development of diabetic complications. In this study,
               AGEs-related pentosidine (PENT) crosslinks-induced structural and
               biochemical changes are studied using multimodal multiphoton
               imaging, Raman spectroscopy and atomic force microscopy (AFM).
               Decellularized equine pericardium (EP) was glycated with four
               ribose concentrations ranging between 5 and 200 mM and monitored
               for up to 30 days. Two-photon excited fluorescence (TPEF) and
               second harmonic generation (SHG) microscopic imaging probed
               elastin and collagen fibers, respectively. The glycated EP showed
               a decrease in the SHG intensities associated with loss of
               non-centrosymmetry of collagen and an increase of TPEF
               intensities associated with PENT crosslinks upon glycation. TPEF
               signals from elastin fibers were unaffected. A three-dimensional
               reconstruction with SHG + TPEF z-stack images visualized the
               distribution of collagen and elastin within the EP volume matrix.
               In addition, Raman spectroscopy (RS) detected changes in
               collagen-related bands and discriminated glycated from untreated
               EP. Furthermore, AFM scans showed that the roughness increases
               and the D-unit structure of fibers remained unchanged during
               glycation. The PENT crosslinked-induced changes are discussed in
               the context of previous studies of glutaraldehyde- and
               genipin-induced crosslinking and collagenase-induced digestion of
               collagen. We conclude that TPEF, SHG, RS, and AFM are effective,
               label-free, and non-destructive methods to investigate glycated
               tissues, differentiate crosslinking processes, and characterize
               general collagen-associated and disease-related changes, in
               particular by their RS fingerprints.",
  month     =  oct,
  year      =  2023,
  keywords  = "Advanced glycation end products; Atomic force microscopy;
               Collagen crosslinking; Raman spectroscopy; Second harmonic
               generation; Two-photon excited fluorescence",
  language  = "en"
}

@ARTICLE{Mostaco-Guidolin2017-ir,
  title     = "Imaging collagen in scar tissue: Developments in second harmonic
               generation microscopy for biomedical applications",
  author    = "Mostaço-Guidolin, Leila and Rosin, Nicole L and Hackett,
               Tillie-Louise",
  journal   = "Int. J. Mol. Sci.",
  publisher = "MDPI AG",
  volume    =  18,
  number    =  8,
  pages     =  1772,
  abstract  = "The ability to respond to injury with tissue repair is a
               fundamental property of all multicellular organisms. The
               extracellular matrix (ECM), composed of fibrillar collagens as
               well as a number of other components is dis-regulated during
               repair in many organs. In many tissues, scaring results when the
               balance is lost between ECM synthesis and degradation.
               Investigating what disrupts this balance and what effect this can
               have on tissue function remains an active area of research.
               Recent advances in the imaging of fibrillar collagen using second
               harmonic generation (SHG) imaging have proven useful in enhancing
               our understanding of the supramolecular changes that occur during
               scar formation and disease progression. Here, we review the
               physical properties of SHG, and the current nonlinear optical
               microscopy imaging (NLOM) systems that are used for SHG imaging.
               We provide an extensive review of studies that have used SHG in
               skin, lung, cardiovascular, tendon and ligaments, and eye tissue
               to understand alterations in fibrillar collagens in scar tissue.
               Lastly, we review the current methods of image analysis that are
               used to extract important information about the role of fibrillar
               collagens in scar formation.",
  month     =  aug,
  year      =  2017,
  keywords  = "fibrillar collagen; image analysis; lung; nonlinear optical
               microscopy; scar tissue; second harmonic generation; skin;
               vessels",
  language  = "en"
}

@ARTICLE{Hristu2025-hh,
  title     = "Quantitative imaging of collagen fibrils via correlated far-field
               and near-field optical techniques",
  author    = "Hristu, Radu and Tranca, Denis E and Stanciu, Stefan G and
               Eftimie, Lucian G and Stanciu, George A",
  journal   = "ACS Mater. Lett.",
  publisher = "American Chemical Society (ACS)",
  pages     = "3642--3651",
  abstract  = "Collagen fibrils are essential for the structural and functional
               integrity of biological tissues, and their imaging is key to
               understanding tissue health and pathology. We present a
               multimodal workflow combining far-field polarization-resolved
               second harmonic generation (PSHG) microscopy with scattering-type
               scanning near-field optical microscopy (s-SNOM) for comprehensive
               collagen characterization. PSHG provides diffraction-limited
               information on fibril anisotropy and helical pitch angle, while
               s-SNOM, paired with atomic force microscopy, yields nanoscale
               optical properties such as refractive index and extinction
               coefficient alongside topographic context. Using type I collagen
               fibrils from bovine tendons, we examine structural and optical
               changes induced by noncollagenous matrix elements, laser-induced
               denaturation, oxidative stress, and thermal exposure. These
               experiments demonstrate the versatility of our approach in
               probing subtle collagen structural details and reveal
               complementary insights into collagen degradation beyond the reach
               of conventional far-field techniques.",
  month     =  oct,
  year      =  2025,
  language  = "en"
}

@INCOLLECTION{Bueno2016-mq,
  title     = "Second harmonic generation microscopy: A tool for quantitative
               analysis of tissues",
  author    = "Bueno, Juan M and Ávila, Francisco J and Artal, Pablo",
  editor    = "Stanciu, Stefan G",
  booktitle = "Microscopy and Analysis",
  publisher = "InTech",
  abstract  = "Second harmonic generation (SHG) is a second‐order non‐linear
               optical process produced in birefringent crystals or in
               biological tissues with non‐centrosymmetric structure such as
               collagen or microtubules structures. SHG signal originates from
               two excitation photons which interact with the material and are
               “reconverted” to form a new emitted photon with half of
               wavelength. Although theoretically predicted by Maria
               Göpert‐Mayer in 1930s, the experimental SHG demonstration arrived
               with the invention of the laser in the 1960s. SHG was first
               obtained in ruby by using a high excitation oscillator. After
               that starting point, the harmonic generation reached an
               increasing interest and importance, based on its applications to
               characterize biological tissues using multiphoton microscopes. In
               particular, collagen has been one of the most often analyzed
               structures since it provides an efficient SHG signal. In late
               1970s, it was discovered that SHG signal took place in
               three‐dimensional optical interaction at the focal point of a
               microscope objective with high numerical aperture. This finding
               allowed researchers to develop microscopes with 3D submicron
               resolution and an in depth analysis of biological specimens.
               Since SHG is a polarization‐sensitive non‐linear optical process,
               the implementation of polarization into multiphoton microscopes
               has allowed the study of both molecular architecture and fibrilar
               distribution of type‐I collagen fibers. The analysis of
               collagen‐based structures is particularly interesting since they
               represent 80\% of the connective tissue of the human body. On the
               other hand, more recent techniques such as pulse compression of
               laser pulses or adaptive optics have been applied to SHG
               microscopy in order to improve the visualization of features. The
               combination of these techniques permit the reduction of the laser
               power required to produce efficient SHG signal and therefore
               photo‐toxicity and photo‐damage are avoided (critical parameters
               in biomedical applications). Some pathologies such as cancer or
               fibrosis are related to collagen disorders. These are thought to
               appear at molecular scale before the micrometric structure is
               affected. In this sense, SHG imaging has emerged as a powerful
               tool in biomedicine and it might serve as a non‐invasive early
               diagnosis technique. © 2016 The Author(s). Licensee InTech. This
               chapter is distributed under the terms of the Creative Commons
               Attribution License (http://creativecommons.org/licenses/by/3.0),
               which permits unrestricted use, distribution, and reproduction in
               any medium, provided the original work is properly cited.",
  month     =  sep,
  year      =  2016
}

@ARTICLE{Tits2021-jk,
  title     = "Joining soft tissues to bone: Insights from modeling and
               simulations",
  author    = "Tits, Alexandra and Ruffoni, Davide",
  journal   = "Bone Rep.",
  publisher = "Elsevier BV",
  volume    =  14,
  number    =  100742,
  pages     =  100742,
  abstract  = "Entheses are complex multi-tissue regions of the musculoskeletal
               system serving the challenging task of connecting highly
               dissimilar materials such as the compliant tendon to the much
               stiffer bone, over a very small region. The first aim of this
               review is to highlight mathematical and computational models that
               have been developed to investigate the many attachment strategies
               present at entheses at different length scales. Entheses are also
               relevant in the medical context due to the high prevalence of
               orthopedic injuries requiring the reattachment of tendons or
               ligaments to bone, which are associated with a rather poor
               long-term clinical outcome. The second aim of the review is to
               report on the computational works analyzing the whole tendon to
               bone complex as well as targeting orthopedic relevant issues.
               Modeling approaches have provided important insights on anchoring
               mechanisms and surgical repair strategies, that would not have
               been revealed with experiments alone. We intend to demonstrate
               the necessity of including, in future models, an enriched
               description of enthesis biomechanical behavior in order to
               unravel additional mechanical cues underlying the development,
               the functioning and the maintaining of such a complex biological
               interface as well as to enhance the development of novel
               biomimetic adhesive, attachment procedures or tissue engineered
               implants.",
  month     =  jun,
  year      =  2021,
  keywords  = "Bimaterial attachment; Bone-tendon interface; Computer
               simulations; Enthesis; Fibrocartilage; Finite element analysis;
               Gradients; Modeling",
  language  = "en"
}

@ARTICLE{Benjamin2002-iz,
  title     = "The skeletal attachment of tendons--tendon ``entheses''",
  author    = "Benjamin, M and Kumai, T and Milz, S and Boszczyk, B M and
               Boszczyk, A A and Ralphs, J R",
  journal   = "Comp. Biochem. Physiol. A Mol. Integr. Physiol.",
  publisher = "Elsevier BV",
  volume    =  133,
  number    =  4,
  pages     = "931--945",
  abstract  = "Tendon entheses can be classed as fibrous or fibrocartilaginous
               according to the tissue present at the skeletal attachment site.
               The former can be ``bony'' or ``periosteal'', depending on
               whether the tendon is directly attached to bone or indirectly to
               it via the periosteum. At fibrocartilaginous entheses, the
               uncalcified fibrocartilage dissipates collagen fibre bending and
               tendon narrowing away from the tidemark; calcified fibrocartilage
               anchors the tendon to the bone and creates a diffusion barrier
               between the two. Where there are additional fibrocartilaginous
               specialisations in the tendon and/or bone next to the enthesis,
               an ``enthesis organ'' is created that reduces wear and tear.
               Little attention has been paid to bone at entheses, despite the
               obvious bearing this has on the mechanical properties of the
               interface and the clinical importance of avulsion fractures.
               Disorders at entheses (enthesopathies) are common and occur in
               conditions such as diffuse idiopathic skeletal hyperostosis and
               the seronegative spondyloarthropathies. They are also commonly
               seen as sporting injuries such as tennis elbow and jumper's knee.",
  month     =  dec,
  year      =  2002,
  language  = "en"
}

@ARTICLE{Stylianou2019-al,
  title     = "Atomic force microscopy on biological materials related to
               pathological conditions",
  author    = "Stylianou, Andreas and Kontomaris, Stylianos-Vasileios and Grant,
               Colin and Alexandratou, Eleni",
  journal   = "Scanning",
  publisher = "Hindawi Limited",
  volume    =  2019,
  pages     =  8452851,
  abstract  = "Atomic force microscopy (AFM) is an easy-to-use, powerful,
               high-resolution microscope that allows the user to image any
               surface and under any aqueous condition. AFM has been used in the
               investigation of the structural and mechanical properties of a
               wide range of biological matters including biomolecules,
               biomaterials, cells, and tissues. It provides the capacity to
               acquire high-resolution images of biosamples at the nanoscale and
               allows at readily carrying out mechanical characterization. The
               capacity of AFM to image and interact with surfaces, under
               physiologically relevant conditions, is of great importance for
               realistic and accurate medical and pharmaceutical applications.
               The aim of this paper is to review recent trends of the use of
               AFM on biological materials related to health and sickness.
               First, we present AFM components and its different imaging modes
               and we continue with combined imaging and coupled AFM systems.
               Then, we discuss the use of AFM to nanocharacterize collagen, the
               major fibrous protein of the human body, which has been
               correlated with many pathological conditions. In the next
               section, AFM nanolevel surface characterization as a tool to
               detect possible pathological conditions such as osteoarthritis
               and cancer is presented. Finally, we demonstrate the use of AFM
               for studying other pathological conditions, such as Alzheimer's
               disease and human immunodeficiency virus (HIV), through the
               investigation of amyloid fibrils and viruses, respectively.
               Consequently, AFM stands out as the ideal research instrument for
               exploring the detection of pathological conditions even at very
               early stages, making it very attractive in the area of bio- and
               nanomedicine.",
  month     =  may,
  year      =  2019,
  language  = "en"
}

@ARTICLE{Bai2023-qo,
  title     = "Deep learning-enabled virtual histological staining of biological
               samples",
  author    = "Bai, Bijie and Yang, Xilin and Li, Yuzhu and Zhang, Yijie and
               Pillar, Nir and Ozcan, Aydogan",
  journal   = "Light Sci. Appl.",
  publisher = "Nature Publishing Group",
  volume    =  12,
  number    =  1,
  pages     =  57,
  abstract  = "Histological staining is the gold standard for tissue examination
               in clinical pathology and life-science research, which visualizes
               the tissue and cellular structures using chromatic dyes or
               fluorescence labels to aid the microscopic assessment of tissue.
               However, the current histological staining workflow requires
               tedious sample preparation steps, specialized laboratory
               infrastructure, and trained histotechnologists, making it
               expensive, time-consuming, and not accessible in resource-limited
               settings. Deep learning techniques created new opportunities to
               revolutionize staining methods by digitally generating
               histological stains using trained neural networks, providing
               rapid, cost-effective, and accurate alternatives to standard
               chemical staining methods. These techniques, broadly referred to
               as virtual staining, were extensively explored by multiple
               research groups and demonstrated to be successful in generating
               various types of histological stains from label-free microscopic
               images of unstained samples; similar approaches were also used
               for transforming images of an already stained tissue sample into
               another type of stain, performing virtual stain-to-stain
               transformations. In this Review, we provide a comprehensive
               overview of the recent research advances in deep learning-enabled
               virtual histological staining techniques. The basic concepts and
               the typical workflow of virtual staining are introduced, followed
               by a discussion of representative works and their technical
               innovations. We also share our perspectives on the future of this
               emerging field, aiming to inspire readers from diverse scientific
               fields to further expand the scope of deep learning-enabled
               virtual histological staining techniques and their applications.",
  month     =  mar,
  year      =  2023,
  language  = "en"
}

@ARTICLE{Madusanka2023-nc,
  title     = "Impact of {H\&E} stain normalization on deep learning models in
               cancer image classification: Performance, complexity, and
               trade-offs",
  author    = "Madusanka, Nuwan and Jayalath, Pramudini and Fernando, Dileepa
               and Yasakethu, Lasith and Lee, Byeong-Il",
  journal   = "Cancers (Basel)",
  publisher = "MDPI AG",
  volume    =  15,
  number    =  16,
  pages     =  4144,
  abstract  = "Accurate classification of cancer images plays a crucial role in
               diagnosis and treatment planning. Deep learning (DL) models have
               shown promise in achieving high accuracy, but their performance
               can be influenced by variations in Hematoxylin and Eosin (H\&E)
               staining techniques. In this study, we investigate the impact of
               H\&E stain normalization on the performance of DL models in
               cancer image classification. We evaluate the performance of
               VGG19, VGG16, ResNet50, MobileNet, Xception, and InceptionV3 on a
               dataset of H\&E-stained cancer images. Our findings reveal that
               while VGG16 exhibits strong performance, VGG19 and ResNet50
               demonstrate limitations in this context. Notably, stain
               normalization techniques significantly improve the performance of
               less complex models such as MobileNet and Xception. These models
               emerge as competitive alternatives with lower computational
               complexity and resource requirements and high computational
               efficiency. The results highlight the importance of optimizing
               less complex models through stain normalization to achieve
               accurate and reliable cancer image classification. This research
               holds tremendous potential for advancing the development of
               computationally efficient cancer classification systems,
               ultimately benefiting cancer diagnosis and treatment.",
  month     =  aug,
  year      =  2023,
  keywords  = "cancer classification; computational complexity; deep learning;
               image classification; stain normalization",
  language  = "en"
}

@ARTICLE{Szulczewski2024-vj,
  title    = "To label or not: the need for validation in label-free imaging",
  author   = "Szulczewski, Joseph M and Yesilkoy, Filiz and Ulland, Tyler K and
              Bartels, Randy and Millis, Bryan A and Boppart, Stephen A and
              Eliceiri, Kevin W",
  journal  = "J. Biomed. Opt.",
  volume   =  29,
  number   = "Suppl 2",
  pages    = "S22717",
  abstract = "Significance: Advances in label-free imaging have impacted many
              areas of biological and biomedical imaging ranging from cell
              biology and cancer to pathology and neuroscience. Despite the
              great progress and advantages of these methods, it is clear that
              to realize their full potential, validation by extrinsic labels
              and probes is critically needed. Aim: This perspective calls for
              developing and applying innovative labels and probes to validate
              both existing and emerging label-free imaging methods. Approach:
              Major representative types of label-free imaging methods are
              briefly presented discussing their advantages and differing
              contrasts. Their biological applications are also reviewed with a
              focus on how validation of label-free methods with carefully
              developed labeling approaches will greatly aid in further
              intrinsic contrast imaging adoption and likely lead to more
              sophisticated image-based biomarkers and a better understanding of
              the underlying signals. Conclusions: Expanded efforts in extrinsic
              label validation will significantly push forward the utilization
              and adoption of label-free methods both in basic research and
              clinical models.",
  month    =  jun,
  year     =  2024,
  keywords = "autofluorescence; intrinsic contrast; label-free imaging; probes;
              validation",
  language = "en"
}

@ARTICLE{Ghosh2023-hb,
  title     = "Viewing life without labels under optical microscopes",
  author    = "Ghosh, Biswajoy and Agarwal, Krishna",
  journal   = "Commun. Biol.",
  publisher = "Nature Publishing Group",
  volume    =  6,
  number    =  1,
  pages     =  559,
  abstract  = "Optical microscopes today have pushed the limits of speed,
               quality, and observable space in biological specimens
               revolutionizing how we view life today. Further, specific
               labeling of samples for imaging has provided insight into how
               life functions. This enabled label-based microscopy to percolate
               and integrate into mainstream life science research. However, the
               use of labelfree microscopy has been mostly limited, resulting in
               testing for bio-application but not bio-integration. To enable
               bio-integration, such microscopes need to be evaluated for their
               timeliness to answer biological questions uniquely and establish
               a long-term growth prospect. The article presents key label-free
               optical microscopes and discusses their integrative potential in
               life science research for the unperturbed analysis of biological
               samples.",
  month     =  may,
  year      =  2023,
  language  = "en"
}

@ARTICLE{Kilpatrick2018-ve,
  title     = "Nanomechanics of cells and biomaterials studied by atomic force
               microscopy",
  author    = "Kilpatrick, Jason I and Revenko, Irène and Rodriguez, Brian J",
  journal   = "Adv. Healthc. Mater.",
  publisher = "Wiley",
  volume    =  7,
  number    =  21,
  pages     = "e1801142",
  month     =  nov,
  year      =  2018,
  language  = "en"
}

@ARTICLE{Zhou2022-kd,
  title     = "Atomic force microscopy ({AFM}) and its applications to
               bone-related research",
  author    = "Zhou, Yuxiao and Du, Jing",
  journal   = "Prog. Biophys. Mol. Biol.",
  publisher = "Elsevier BV",
  volume    =  176,
  pages     = "52--66",
  abstract  = "Bone consists of organic (mostly collagen) and inorganic (mostly
               bioapatite mineral) components that are organized into
               hierarchical structures from nano-to macro-scales that provide
               load-bearing functions. The structures and properties of bone are
               affected by bone remodeling activities, which are affected by
               mechanotransduction, a process through which mechanical signals
               are converted to biochemical signals in cellular signaling.
               Atomic Force Microscopy (AFM) technique can be used to
               characterize the surface morphology and mechanical properties of
               the specimens and can achieve atomic resolution in the resulting
               images. Therefore, the AFM technique has been applied in bone
               research and has provided new understandings of the structures
               and properties of hierarchical structures in bone across multiple
               length scales. This review begins by introducing the tip-surface
               interactions and the operation modes of AFM, including the
               recently developed sub-resonance modes, including PeakForce
               Tapping mode. Then the contact adhesion theories used in
               analyzing AFM data are reviewed, followed by a systematic review
               of the applications of the AFM technique to bone and bone-related
               tissues and cells, including surface morphology imaging, contact
               indentation testing, and other mechanical tests. The applications
               of sub-resonance tapping mode to bone and other biological
               molecules, cells, and tissues are also reviewed.",
  month     =  dec,
  year      =  2022,
  keywords  = "Atomic force microscopy (AFM); Bone; Mechanical properties;
               Sub-resonance tapping; Surface morphology",
  language  = "en"
}

@ARTICLE{Shioka2024-ao,
  title     = "Ex vivo {SIM}-{AFM} measurements reveal the spatial correlation
               of stiffness and molecular distributions in {3D} living tissue",
  author    = "Shioka, Itsuki and Morita, Ritsuko and Yagasaki, Rei and
               Wuergezhen, Duligengaowa and Yamashita, Tadahiro and Fujiwara,
               Hironobu and Okuda, Satoru",
  journal   = "Acta Biomater.",
  publisher = "Elsevier BV",
  volume    =  189,
  pages     = "351--365",
  abstract  = "Living tissues each exhibit a distinct stiffness, which provides
               cells with key environmental cues that regulate their behaviors.
               Despite this significance, our understanding of the
               spatiotemporal dynamics and the biological roles of stiffness in
               three-dimensional tissues is currently limited due to a lack of
               appropriate measurement techniques. To address this issue, we
               propose a new method combining upright structured illumination
               microscopy (USIM) and atomic force microscopy (AFM) to obtain
               precisely coordinated stiffness maps and biomolecular
               fluorescence images of thick living tissue slices. Using mouse
               embryonic and adult skin as a representative tissue with
               mechanically heterogeneous structures inside, we validate the
               measurement principle of USIM-AFM. Live measurement of tissue
               stiffness distributions revealed the highly heterogeneous
               mechanical nature of skin, including nucleated/enucleated
               epithelium, mesenchyme, and hair follicle, as well as the role of
               collagens in maintaining its integrity. Furthermore, quantitative
               analysis comparing stiffness distributions in live tissue samples
               with those in preserved tissues, including formalin-fixed and
               cryopreserved tissue samples, unveiled the distinct impacts of
               preservation processes on tissue stiffness patterns. This series
               of experiments highlights the importance of live mechanical
               testing of tissue-scale samples to accurately capture the true
               spatiotemporal variations in mechanical properties. Our USIM-AFM
               technique provides a new methodology to reveal the dynamic nature
               of tissue stiffness and its correlation with biomolecular
               distributions in live tissues and thus could serve as a technical
               basis for exploring tissue-scale mechanobiology. STATEMENT OF
               SIGNIFICANCE: Stiffness, a simple mechanical parameter, has drawn
               attention in understanding the mechanobiological principles
               underlying the homeostasis and pathology of living tissues. To
               explore tissue-scale mechanobiology, we propose a technique
               integrating an upright structured illumination microscope and an
               atomic force microscope. This technique enables live measurements
               of stiffness distribution and fluorescent observation of thick
               living tissue slices. Experiments revealed the highly
               heterogeneous mechanical nature of mouse embryonic and adult skin
               in three dimensions and the previously unnoticed influences of
               preservation techniques on the mechanical properties of tissue at
               microscopic resolution. This study provides a new technical
               platform for live stiffness measurement and biomolecular
               observation of tissue-scale samples with micron-scale resolution,
               thus contributing to future studies of tissue- and organ-scale
               mechanobiology.",
  month     =  nov,
  year      =  2024,
  keywords  = "Atomic force microscopy; Elastic modulus; Embryonic skin; Ex vivo
               culture; Soft tissue mechanics",
  language  = "en"
}
