## Supplementary material for "Correlative SHG-AFM imaging workflow for label-free quantitative analysis of collagen structure–function relationships": LaTEX: sn-MAIN-article.tex

%Version 3.1 December 2024
% See section 11 of the User Manual for version history
%
%%%%%%%%%%%%%%%%%%%%%%%%%%%%%%%%%%%%%%%%%%%%%%%%%%%%%%%%%%%%%%%%%%%%%%
%%                                                                 %%
%% Please do not use \input{...} to include other tex files.       %%
%% Submit your LaTeX manuscript as one .tex document.              %%
%%                                                                 %%
%% All additional figures and files should be attached             %%
%% separately and not embedded in the \TeX\ document itself.       %%
%%                                                                 %%
%%%%%%%%%%%%%%%%%%%%%%%%%%%%%%%%%%%%%%%%%%%%%%%%%%%%%%%%%%%%%%%%%%%%%

%%\documentclass[referee,sn-basic]{sn-jnl}% referee option is meant for double line spacing

%%=======================================================%%
%% to print line numbers in the margin use lineno option %%
%%=======================================================%%

%%\documentclass[lineno,pdflatex,sn-basic]{sn-jnl}% Basic Springer Nature Reference Style/Chemistry Reference Style

%%=========================================================================================%%
%% the documentclass is set to pdflatex as default. You can delete it if not appropriate.  %%
%%=========================================================================================%%

%%\documentclass[sn-basic]{sn-jnl}% Basic Springer Nature Reference Style/Chemistry Reference Style

%%Note: the following reference styles support Namedate and Numbered referencing. By default the style follows the most common style. To switch between the options you can add or remove Numbered in the optional parenthesis. 
%%The option is available for: sn-basic.bst, sn-chicago.bst%  
 
\documentclass[pdflatex,sn-nature]{sn-jnl}% Style for submissions to Nature Portfolio journals
%\documentclass[pdflatex,sn-basic]{sn-jnl}% Basic Springer Nature Reference Style/Chemistry Reference Style
%%\documentclass[pdflatex,sn-mathphys-num]{sn-jnl}% Math and Physical Sciences Numbered Reference Style
%%\documentclass[pdflatex,sn-mathphys-ay]{sn-jnl}% Math and Physical Sciences Author Year Reference Style
%%\documentclass[pdflatex,sn-aps]{sn-jnl}% American Physical Society (APS) Reference Style
%%\documentclass[pdflatex,sn-vancouver-num]{sn-jnl}% Vancouver Numbered Reference Style
%%\documentclass[pdflatex,sn-vancouver-ay]{sn-jnl}% Vancouver Author Year Reference Style
%%\documentclass[pdflatex,sn-apa]{sn-jnl}% APA Reference Style
%%\documentclass[pdflatex,sn-chicago]{sn-jnl}% Chicago-based Humanities Reference Style

%%%% Standard Packages
%%<additional latex packages if required can be included here>

\usepackage{graphicx}%
\usepackage{multirow}%
\usepackage{amsmath,amssymb,amsfonts}%
\usepackage{amsthm}%
\usepackage{mathrsfs}%
\usepackage[title]{appendix}%
\usepackage{xcolor}%
\usepackage{textcomp}%
\usepackage{manyfoot}%
\usepackage{booktabs}%
\usepackage{algorithm}%
\usepackage{algorithmicx}%
\usepackage{algpseudocode}%
\usepackage{listings}%
\usepackage{gensymb}%
%% Added packages - HM
\usepackage{svg}
\usepackage{wrapfig}
%%%%

%%%%%=============================================================================%%%%
%%%%  Remarks: This template is provided to aid authors with the preparation
%%%%  of original research articles intended for submission to journals published 
%%%%  by Springer Nature. The guidance has been prepared in partnership with 
%%%%  production teams to conform to Springer Nature technical requirements. 
%%%%  Editorial and presentation requirements differ among journal portfolios and 
%%%%  research disciplines. You may find sections in this template are irrelevant 
%%%%  to your work and are empowered to omit any such section if allowed by the 
%%%%  journal you intend to submit to. The submission guidelines and policies 
%%%%  of the journal take precedence. A detailed User Manual is available in the 
%%%%  template package for technical guidance.
%%%%%=============================================================================%%%%

%% as per the requirement new theorem styles can be included as shown below
\theoremstyle{thmstyleone}%
\newtheorem{theorem}{Theorem}%  meant for continuous numbers
%%\newtheorem{theorem}{Theorem}[section]% meant for sectionwise numbers
%% optional argument [theorem] produces theorem numbering sequence instead of independent numbers for Proposition
\newtheorem{proposition}[theorem]{Proposition}% 
%%\newtheorem{proposition}{Proposition}% to get separate numbers for theorem and proposition etc.

\theoremstyle{thmstyletwo}%
\newtheorem{example}{Example}%
\newtheorem{remark}{Remark}%
\theoremstyle{thmstylethree}%
\newtheorem{definition}{Definition}%
\raggedbottom
\unnumbered% uncomment this for unnumbered level heads
\setcounter{secnumdepth}{2} % adds nubmers to subsections
\setcitestyle{super,comma,sort&compress}

\begin{document}

\title[Article Title]{Correlative SHG-AFM imaging workflow for label-free quantitative analysis of collagen structure-function relationships}

\equalcont{These authors contributed equally to this work.}

\equalcont{These authors contributed equally to this work.}






\affil[1]{\orgdiv{California NanoSystems Institute}, \orgname{University of California Los Angeles}, \orgaddress{ \city{Los Angeles}, \state{CA}, \country{USA}}}

\affil[2]{\orgdiv{Department of Orthopedic Surgery}, \orgname{University of California Los Angeles}, \orgaddress{ \city{Los Angeles}, \state{CA}, \country{USA}}}


%%==================================%%
%% Sample for unstructured abstract %%
%%==================================%%

\abstract{We present a user-friendly correlative second harmonic generation (SHG) and atomic force microscopy (AFM) imaging workflow for quantifying the nanomechanical properties of collagen in unfixed, unlabeled tissue sections. SHG Aligned Profiling for Elasticity and Segmentation or SHAPES utilizes SHG imaging to guide AFM force mapping, enabling label-free, anatomically specific selection of regions of interest, facilitating spatially resolved characterization of fibrous collagen morphology and local stiffness. Compared to brightfield or confocal contrast-guided AFM mapping, SHG improves anatomical specificity without fixation or staining, enabling downstream analysis on the same tissue section while preserving spatial correspondence between structural (SHG) and mechanical (AFM) data. Rapid identification of regions of interest also reduces the risk of costly AFM tip breakage, improving throughput and reducing operator burden. Utilizing only standard turn-key commercial systems common in many user facilities and clinical laboratories, multimodal image coregistration and automated identification of anatomical regions of interest are employed to integrate SHG and AFM datasets across complex sample topographies. Coregistration of SHG and AFM images substantially increases the number of usable datasets per measurement session, facilitating translation of these complementary modalities and bridging nanomechanical imaging with clinical research practice.

 }
\keywords{collagen, second harmonic generation, two photon excitation microscopy, multiphoton, atomic force microscopy, correlative imaging}


\maketitle

\section{Introduction}\label{sec1}
Characterization and analysis of biologic tissues by light microscopy has historically relied on exogenous staining to identify and classify structural features. Collagen is a principal structural component of the extracellular matrix (ECM) and a major determinant of tissue architecture, mechanics, and function across a wide range of organ systems. As alterations in collagen fiber assembly, orientation, and crosslinking are central to the pathogenesis of fibrosis, cancer, connective tissue disorders, vascular disease, musculoskeletal degeneration, abnormal wound healing, and many other pathologies, collagen is one of the most widely targeted sources of contrast in light microscopy. \cite{Mostaco-Guidolin2017-ir},\cite{James2021-zt} The standard stain for collagen is picrosirius red (PSR),\cite{Liu2017-wm} a polarization sensitive stain which amplifies the natural birefringence of collagens type I/II by binding preferentially and coherently to the already anisotropic fibrous collagen structure.\cite{Lopez-De-Padilla2021-hb} However, current methods generally rely on tissue fixation that generates artifacts and adds variability to the staining results. Despite the ubiquity of tissue stains and fluorescent probes, many of these methods still require validation in certain applications.\cite{Szulczewski2024-vj} With the emergence of virtual stains\cite{Bai2023-qo} and automated segmentation using machine learning (ML) the use of stained images as reference data sets has exposed limitations due to the introduction of variations in appearance between tissue types, protocols used, and reproducibility from different laboratories.\cite{Madusanka2023-nc} Machine learning based tissue identification is highly dependent on the quality and reproducibility of the images used for model training, and there is an increasing push towards label-free microscopy techniques as an alternative.\cite{Ghosh2023-hb} In the past twenty years Second Harmonic Generation (SHG) imaging has increasingly been used as a label-free alternative to collagen imaging,\cite{Williams2005-id,Cisek2021-fy,Mostaco-Guidolin2017-ir} with the additional benefit of enabling deeper penetration depths \cite{Crosignani2014-do} in large centimeter sized bulk surgical tissue samples.\cite{Kashani-Ligumsky2026-nt} The incorporation of SHG imaging to map fibrillar collagen without staining or fixation allows mapping of one of the most mechanically significant tissue components compatible with quantitative mechanical measurements of the same section enabling rigorous assessment of structure-function relationships which often rely solely on inference from structure \cite{Plekhanov2023-yb, Arshee2025-wg}. 

Atomic force microscopy (AFM) is increasingly utilized to produce high resolution topographic and mechanical maps of biological samples that provide distinct information without relying on artificial staining or optical labeling. For the study of tissue mechanics, this modality generates quantifiable metrics of elasticity across large 2D regions that can be combined with light microscopy images to determine the relationship between structural and mechanical features of tissues. Early \cite{Plodinec2010-wg} applications of AFM mechanical imaging to articular cartilage tissue enabled the quantification of the Young's modulus across the surface of osteochondral plugs and differentiated the stiffness of collagen fibers from the softer extracellular matrix. Subsequent extension of AFM-based tissue mechanics was limited for many years by the design of the microscope itself, which historically focused on measuring extremely smooth, flat samples, optimized for atomic-scale precision at the expense of scanning range and sample roughness. Recent developments have extended the useful range of the AFM scanner in the vertical (Z) direction from the typical 10-15 \(\mu\)m to over 200 \(\mu\)m across a large scan area (X/Y). A combination of high-accuracy motorized stages and seamless integration with optical stereomicroscopes delivers access to regions over centimeter length scales, facilitating mechanical mapping of cryotome and vibratome prepared tissue samples even when samples features vary more than 10-20 \(\mu\)m between adjacent pixels. These technology improvements make AFM a practical, compatible imaging modality for common clinical samples such as cryotome sliced sections that are routinely used in histology \cite{Stylianou2019-al, Cho2023-rt}. 

The enthesis, the tendon-to-bone insertion site, is of major interest to musculoskeletal researchers and a key application area for both multiphoton and AFM microscopy, as it is inherently difficult to image using conventional staining approaches due to its steep compositional and structural gradients spanning soft collagenous tendon, proteoglycan-rich fibrocartilage, and highly mineralized bone over micrometer length scales.\cite{de-Palma2004-by,Tits2021-jk} These transitions produce significant variability in stain uptake, light scattering, and optical contrast, limiting the reliability of boundary identification in standard histological preparations \cite{Silva2022-dy}. Furthermore, decalcification protocols required for mineralized tissue introduce structural distortions and can disrupt collagen architecture, while fixation alters mechanical properties, complicating correlative structure–function analyses and requiring laborious protocols for satisfactory results. \cite{Angrisani2022-eh}

While the independent utility of both SHG/2PE and AFM in characterizing vascular tissue and mechanics has been demonstrated\cite{Li2025-wb}, these modalities have not been deployed in a formally correlative framework. This has resulted in insufficient characterization of structure-function relationships in many tissue types. We present SHAPES (SHG Aligned Profiling for Elasticity and Segmentation), a label-free correlative imaging workflow combining SHG and atomic force microscopy to quantitatively assess collagen microstructure and local stiffness in native tissues (Fig~\ref{schema}). The four-step workflow: cryosectioning, multiphoton imaging for ROI selection, AFM mechanical mapping, and deep learning enabled image segmentation can be carried out on standard commercial instruments without the need for specialized hardware or custom software. SHAPES is fully compatible with downstream fixation, staining, and image analysis pipelines \cite{Szulczewski2024-vj}, and captures two photon autofluorescence (2PE) with optional fluorescence lifetime imaging (FLIM), generating label-free contrast data to further assist in locating the demarcation lines of enthesis tissues. The method is compatible with fresh whole mounted biopsies, vibratome-sliced living tissue, and cryogenically preserved sections that avoid fixation which would modify the interpretation of mechanical data \cite{Plodinec2010-wg}. Landmark based coregistration of SHG/2PE maximum projections with brightfield images from the microscope built into the AFM enables precise spatial correspondence between structural and mechanical datasets without fiducial markers \cite{Hristu2025-hh}. Last, easily being able to not just coregister but also automatically segment full mapped ROIs provides investigators and clinicians with higher-dimensional, more reproducible multimodal characterization than either modality alone \cite{Shioka2024-ao,Quansah2023-um, Jones2022-zk, Li2025-wb}. In this study, we apply the SHAPES protocol to the enthesis and skeletal muscle, demonstrating a label-free workflow that integrates SHG and 2PE autofluorescence to resolve the bone–tendon interface and guide precise correlation with AFM within the same imaging platform. This approach enables targeted elasticity mapping that reveals unexpected mechanical heterogeneities at the enthesis, while machine learning–guided segmentation distinguishes bone, cartilage, and tendon compartments based on their distinct mechanical signatures. Finally, we show that SHAPES generalizes to skeletal muscle, where SHG-guided AFM resolves the endomysium and individual myofiber compartments.

\begin{figure}[h!] 
    \centering
    \includesvg[width=1\linewidth, inkscapelatex=false]{Figures/fig1.svg} 
    \caption{Schema of the SHAPES (SHG aligned profiling of elasticity) workflow: (a) Cryosectioning to prepare flat tissue sections mounted on standard microscope slides. (b) SHG/2PE microscopy to generate depth and overview maps for ROI selection. (c) AFM probing of the same slide-mounted section to quantify local mechanical properties. (d) Deep learning enabled segmentation of an SHG/AFM overlay image of a cross-sectional myofiber bundle.}
\label{schema}
\end{figure}

\section{Methods}\label{sec2}
\subsection{Animal surgeries and ethical approval declarations} 
Native, unmanipulated Achilles tendon–calcaneus enthesis specimens were harvested from adult mice following euthanasia under approved UCLA Institutional Animal Research Committee protocols and immediately fresh-frozen for cryosectioning to establish normative nanomechanical reference value, as previously described~\cite{Kremen2023-et,Shi2024-az}. All animal procedures were conducted in accordance with approved UCLA Institutional Animal Research Committee protocols.

Human tissue was obtained from typically developing children aged 11-16 years with no history of neuromuscular disorders, enrolled at a tertiary care academic referral center under UCLA Institutional Review Board approval (IRB 21-000350). Written informed consent was obtained from parents or caregivers, and age-appropriate assent was obtained from all participants. Biopsies of the vastus lateralis (VL) were obtained during anterior cruciate ligament reconstruction, corrective osteotomy, or hardware removal. All muscle samples were transported to our laboratory in a specimen container on ice and processed within 2 hours of resection. Biopsies were placed in labeled cassettes, rapidly frozen in liquid nitrogen, and stored at -80\degree~C until use. 

\subsection{Sample Preparation} Harvested tissue was rapidly frozen, embedded in optimal cutting temperature compound (OCT; Tissue-Tek), and cryosectioned at 50$\mu$m thickness onto glass slides using a cryomicrotome (Leica Cryocut 1800 cryostat), allowed to dry at room temperature for 2 hours, and stored at -80\degree C, allowing the workflow to be paused indefinitely prior to downstream analysis. This provides a stable intermediate stage between tissue extraction and analysis, making it particularly useful for high-throughput processing and longitudinal studies. When necessary to select or examine samples, sections were quickly thawed in PBS and evaluated in bright field mode using a stereo-microscope to determine suitability for SHAPES analysis. (Fig.~\ref{mpm}b). Immediately prior to imaging, sections were removed from the freezer and allowed to acclimate at room temperature for 30 minutes before being placed in PBS and transported.

\subsection{Two-Photon Excitation (2PE) and Second Harmonic Generation (SHG) Microscopies}
Multiphoton microscopy (2PE/SHG) was performed on a Leica SP8 MP-DIVE microscope with a Spectra Physics Insight X3+ laser with a 120 femtosecond pulse width. Imaging was performed with an excitation wavelength of 830 nm in order to simultaneously produce second harmonic generation scattering from type I/II fibrous collagens and generate 2PE autofluorescence of intrinsic fluorophores such as  elastin and heme. Energy diagrams of these label-free processes compared to traditional fluorescence excitation is shown in (Fig.~\ref{jablonski}). The resulting image shows the type I collagen SHG signal in green and the autofluorescence of elastin and other natural fluorophores in magenta (Fig.~\ref{mpm}c). 

The enthesis sample discussed herein was imaged with an excitation wavelength of 830 nm at 25.7\% power out of 1.74 W total and the SHG component was collected from 405 nm to 425 nm and the 2PE component was collected from 522 nm to 532 nm (Fig.~\ref{mpm}(c). Images were collected using a HC PL FLUOTAR L 40x/0.60 dry lens and resonant scanner at 8000 Hz with 6 line averages plus 6 frame averages. Images were collected with 1024 x 1024 X-Y pixels, 220 nm per pixel resulting in a physical size of 221.43 \(\mu\)m per image with 12 bit intensity resolution. 24 Z sections were taken with slice thickness of approximately 4 \(\mu\)m. The ROI imaged comprised of 18 tiles which were stitched together in the X and Y dimensions with a 10\% overlap per image in the Leica LASX Software. The resulting stitched image resulted in a physical size of 608.23 \(\mu\)m X, 619.48 \(\mu\)m Y and 92.03 \(\mu\)m Z dimensions, which was then merged to generate a 2D maximum Z-projection with the same X-Y dimensions as the original stitched image. The intensity of each channel was adjusted to optimize the appearance of the SHG and autofluorescence signals and the composite image was exported as a jpeg for transfer to the AFM for correlative alignment. A 2x2 binned image was also exported for coregistration with the final AFM maps in AIVIA (Leica Microsystems) during segmentation as described in section 2.5. The physical dimensions of the image were recorded and the image was uploaded to an online file repository for transfer to the AFM. Additionally, the raw .lif file containing the 2x2 binned image can also be used directly in AIVIA to ensure the pix-mm conversion is not lost. 


\begin{figure}%{wrapfigure}{l}{0.5\linewidth} 
    \includesvg[width=0.5\textwidth, inkscapelatex=false]{Figures/jablonski.svg} 
    \caption{Jablonski energy diagram comparing traditional fluorescence to the multiphoton microscopy modalities used for label-free imaging herein.}
    \label{jablonski}
\end{figure}


\par\noindent


\subsection{AFM Imaging} 
AFM data was acquired using a JPK Nanowizard 4A microscope equipped with a JPK Hybrid Stage allowing 200 x 200 x 200 \(\mu\)m X-Y-Z scan dimensions. This system is mounted on a Leica M205 FCA stereomicroscope with an Andor Zyla sCMOS camera. To identify the approximate region of the SHG/2PE image on the AFM, the SHG/2PE image was imported directly into the Bruker NW4A control software and the physical dimensions of the image are entered into the manual scaling feature on import to match the dimensions to the scale on the AFM. The image was then rotated 90\degree~ then aligned with the Optical Transform option available through the Viewer Window of the JPK SPM Desktop software. This option allows the representation of the optical image at 50\% transparency on top of bright field images acquired by the M205. Precise alignment was achieved by matching shared intrinsic structural tissue landmarks visible in both modalities, including sample edges, distinct morphological features of the tissue, and fiducial markers when present. A video of this process is shown in Supplemental Video 1. In general, tissue edges are useful landmarks to obtain accurate alignment as well. Once alignment was completed in the AFM software, the SHG/2PE image is used to identify the specific feature, in this case the enthesis region, to select an AFM scan region.  


Force Mapping mode was employed with sufficient Z length to ensure a flat force baseline before contacting the sample. Pixel spacing was chosen to capture the size of the biological features of interest in balance with the total acquisition time and the desired field of view. For this sample, with enthesis features as small as 1 \(\mu\)m, an area of 200$\times100\mu$m X-Y with 500x250 pixel resolution and 500x500 nm pixel spacing was selected.  An AFM probe (Bruker RTESPA-525-30) with a spherical tip (30 nm nominal radius of curvature), spring constant of \~ 200 N/m, and calibrated deflection sensitivity of 20.4 nm/V was used to acquire force distance curves at each pixel of the mechanical map. Force curves were fit to produce a Young's modulus value at each pixel using the JPKSPM Data Processing software. Curve fitting employed six standard processing steps: (1) calibration, (2) smoothing, (3) baseline subtraction, (4) contact point identification, (5) determination of vertical tip position, and (6) elasticity fit with minimal to the curve region used in each step. The fit parameters for the Young's modulus included selection of the Hertz/Sneddon model with a spherical tip shape, tip radius of 30 nm, and a Poisson ratio of 0.50. The resulting elasticity fitting was inspected for several hundred curves to validate proper alignment.  After verifying accurate fitting, batch fitting was applied to determine the Young's modulus for each pixel without further filtering or modification.  No cut off value for exclusion of curves was applied.   The final data must be exported as filetype .ome.tiff for import into AIVIA for automated segmentation analysis.\newline

\subsection{Image Registration and Segmentation}
For image registration, segmentation, and analysis, AIVIA v16.0 (Leica Microsystems) was utilized as outlined in Fig \ref{aivia}(a). First, two supplementary Python files may first be loaded into AIVIA as 'Recipes' to enable rotation and rescaling of pixel sizes between the AFM and SHG images, as needed depending on the orientation, degree of overlap, and pixel to mm conversion factor of the images exported from each system with respect to each other. These files are made available by downloading from the online AIVIA Github community at https://github.com/AiviaCommunity/PythonForAivia. The AFM channel is used as the base layer and then the appropriately oriented SHG/2PE channels are imported as overlaid channels to be used for further training and segmentation as desired.  

For automated segmentation based on AFM Young’s modulus maps, the AIVIA pixel classifier tool was trained to differentiate bone, tendon, and cartilage from background. Briefly, the classifier uses a small set of user-annotated regions as training input to estimate, for each pixel, the probability of belonging to a given class. From these annotations, a feature vector is computed for every pixel in the image. These feature vectors are then used to train a random forest classifier, which assigns class labels to previously unseen pixels. To capture structural information across multiple spatial scales, a series of image processing filters are applied using user-defined kernel sizes, enabling detection of both regional (large-scale) and local (fine-scale) features. The trained model generates “confidence maps,” output as additional image channels, where pixel intensities (0–255) represent the probability of membership in each class. Subsequently, the Smart Segmentation module converts these probability maps into discrete object boundaries using a user-defined confidence threshold. The resulting segmented regions are used for quantitative analysis and can be opened easily as tiff files in ImageJ and other open source software. Additional parameters, including separation factor and object size filtering, can be adjusted to refine segmentation and stratify outputs in the final dataset.


\section{Results}\label{sec3}

\begin{figure}[h!]
    \includesvg[width=1\textwidth, inkscapelatex=false]{Figures/fig2a.svg} 
    \caption{\textit{Visualizing the enthesis via collagen and autofluorescence signals.} (a) Cartoon schematic of the enthesis region of interest, located at the interface of soft tendon and hard mineralized bone. (b) Under standard brightfield imaging, it is difficult to distinguish the enthesis interface.  (c) Representative multiphoton emission spectra under 830nm laser excitation reveals a sharp band at 415nm indicative of SHG from collagen fibers, as well as broadband autofluorescence emission from 450-650nm. (d) The resulting multiphoton overlay image used for contrast in subsequent AFM imaging}
    \label{mpm}
\end{figure}

\subsection{Bone-tendon Interface Resolved by Label-free SHG and 2PE Autofluorescence Contrast} The connection of the Achilles tendon to the calcaneus bone in both mice and humans forms an enthesis region, transitioning from tendon to fibrocartilage to bone as shown in Fig~\ref{mpm}(b). Identifying the boundaries of each tissue type can be difficult without the use of fixation and staining, which conflict with the requirements for mechanical characterization of unmodified tissue. Figure~\ref{mpm}(b) shows a brightfield image of a normal enthesis region for this tendon to bone connection after being cryogenically frozen and cryosliced. This sample was thawed and directly imaged in PBS with both the SHG and 2PE channel with the emitted light collected as shown in Figure~\ref{mpm}(c). The resulting image in Fig~\ref{mpm}(d) reveals the collagen rich tissues that span from the tendon into the bone in the green channel from SHG, and the start of the bone at the enthesis in the magenta channel from 2PE emission. This initial label-free image of an unaltered tissue sample provides a feature rich representation of native tissue that can be utilized to rapidly and reproducibly identify regions of interest in many tissue types, including examples to date in our facility of joint and muscle,\cite{Shi2024-az,Kot2024-lo} lung, pancreas, arteries,\cite{Li2025-wb} muscle, vocal cord,\cite{Schlegel2023-gy} placenta,\cite{Kashani-Ligumsky2026-nt} and epiglottal and laryngeal tisses.\cite{Moffatt2026-qh} 

\subsection{Correlation of SHG/2PE and Brightfield within AFM Imaging Platform Enables Precise Targeted Elasticity Mapping} 

\begin{figure}[h!]
    \includesvg[width=1\linewidth, inkscapelatex=false]{Figures/fig3panel-zoom.svg} 
    \caption{\textit{Correlative imaging enables AFM mapping.} Live JPK brightfield image on the AFM system for mapping elasticity shown for wild type mouse calcaneus Achilles tendons (a,d). Multiphoton images for each region (b,e) help to guide the user to specific small ROIs representative of organized and disorganized collagen structure, and the enthesis tidemark, respectively. Alignment of multiphoton max projection overlay image with live JPK brightfield image enables rapid identification of mapping areas for either c) point-by-point based small ROI analysis or for f) long overnight mapping sessions. All scale bars represent 100 microns.}
    \label{align}
\end{figure}

The ability to import virtually any jpeg image with a known scale into the JPK AFM software to align tissue features visible in multiple modalities at or near the region of interest proved to be a critical time saving factor, particularly in terms of maintaining tissue viability throughout the experiment so that samples could be used for further downstream analysis. Example multiphoton images of cryosliced wild type mouse calcaneus Achilles tendons are shown in Fig \ref{align} (b,e), which provide contrast (green) from second harmonic generation and (magenta) two photon autofluorescence to enable rapid identification of collagen structures and provide a practical alignment process when overlaid into the AFM navigation software, as shown in the last panels (c,f). Without this additional contrast, tissue degradation due to time becomes a risk. With SHG/2PE as a guidance to quickly narrow in on the users specific area of interest, maps of the topography and mechanical stiffness (Young's modulus) can then be taken at distinct small ROI locations as shown in Fig \ref{align} (c) or as widespread map areas such as the enthesis region shown in Fig \ref{align} (f). This flexibility highlights the important advances in AFM analysis of tissue samples in comparison to previous systems where entire studies were often limited to small area samples such as those shown in Fig \ref{align} (c) and the ability to identify specific regions of interest across large intact tissue areas for high resolution mapping provides important new opportunities to leverage AFM imaging in medicine and biology

\subsection{AFM Mapping Reveals Unexpected Mechanical Heterogeneities} Using SHG/2PE images to guide identification of the enthesis region, which can be on the order of 1-10 micrometers in mouse tissue, enabled rapid identification of AFM imaging regions of interest compared to brightfield alone. The traditional topographic (height) image shown in Fig~\ref{afm}a provides some contrast between the bone, and tendon areas, however this can vary based on the positioning of the sample on a slide, as well as the relative lifting or sagging of the sample after cryoslicing. As a standalone measurement, it also fails to characterize the important fibrocartilage regions or process bone structures which can be found within the enthesis region.

A major practical advantage of SHG-guided AFM mapping in mineralized tissue is the time saved from the substantial reduction in cantilever/probe breakage. Undecalcified bone presents unpredictable surface asperities arising from cryosectioning artifacts, including regions where the blade has deflected around hard mineral nodules or where bone fragments have lifted above the surrounding tissue plane. Encountering these features during unattended overnight mapping otherwise routinely results in catastrophic probe failure, terminating the scan early and requiring tip replacement at significant cost. The extended Z-range of the JPK NanoWizard 4A (200~\textmu m in X, Y, and Z) allows the system to traverse these topographic discontinuities without tip contact, provided scan parameters are configured with sufficient Z-travel headroom. Pre-acquisition SHG/2PE imaging identifies bone-rich subregions and informs parameter selection accordingly, converting overnight mapping of undecalcified enthesis sections from an unreliable procedure into a reproducible one.

\begin{figure} [h!]
    \includesvg[width=0.8\textwidth, inkscapelatex=false]{Figures/fig4afm.svg} 
    \caption{\textit{AFM elasticity mapping of the tendon–bone enthesis.} Atomic force microscopy mapping of a mouse enthesis region connecting the Achilles tendon and calcaneus bone a) Height map of enthesis compared to b) Young’s modulus image constructed from fitting indentation curves at every pixel. Representative indentation curves showing a lower stiffness (c, blue) and higher stiffness (d, red). Scale bars 10 microns.}
    \label{afm}
\end{figure}

In contrast, fitting the complete force curves generates the map of the Young's modulus shown in Fig~\ref{afm}b, providing rich mechanically based differentiation of the tissue types, and distinctly identifying the tendon, fibrocartilage, and bone regions. In this image, the upper right region is tendon (red), leading to a fibrocartilage region (black)  surrounding a bone process (yellow), which then connects to the calcaneus bone in the lower left (yellow). By coloring the map according to the elastic modulus, both stiffness and textural tissue details emerge that are not clearly identifiable from the height map or the SHG/2PE image used for initial guidance. Fig \ref{afm} provides example force curves from a soft (c), fibrocartilage) and a hard (d),bone) region demonstrating how the contrast in the stiffness of the materials in represented in the AFM data collected. The Hertz model is used to fit the force curve at each pixel to calculate the a single value for the Young's modulus for each point in Fig \ref{afm} (b).

\subsection{Machine Learning–Guided Segmentation Resolves Bone, Cartilage, and Tendon Elasticity at the Enthesis} AFM modulus maps are easily coregistered with multiphoton images in AIVIA by importing the SHG channel (exported as a single-channel \textbackslash{SHG.aivia.tiff} from the Leica \textbackslash{*.lif} file) into a new 16-bit channel overlayed with the 32-bit float AFM image. This yields a two-channel composite (Fig.~\ref{aivia}a) to which registration recipes were applied for spatial alignment using the public 'PythonforAIVIA' Github repository \url{https://github.com/AiviaCommunity/PythonForAivia/tree/master/PythonEnvForAivia/Recipes/TransformImages}

\begin{figure} [h!]
    \includesvg[width=1\textwidth, inkscapelatex=false]{Figures/fig-aivia1-1.svg} 
    \caption{\textit{Tissue-segmented elasticity mapping of the enthesis.} Visualization of elasticity at the enthesis separated by segmented tissue class. a) Multiphoton image with SHG in green, autofluorescence in magenta, and AFM Modulus map in grey, alongside the b) pixel classifier results for classes bone (lavender), cartilage/enthesis (orange), and tendon (yellow). Final heatmaps of Young's Modulus segmented elasticity maps for the c) bone, d) cartilage/enthesis, and e) tendon regions, where all purple to yellow inferno lookup tables range from 0 to 6 GPa. Scale bar is 10 microns.}
    \label{aivia}
\end{figure}


\begin{table}[h!]
\centering
\caption{Spot-based ROI AFM analysisDescriptive statistics and pairwise Wilcoxon rank-sum test results 
(Bonferroni corrected) for Young's modulus measurements across tissue classes }
\label{tab:wilcox_spot}
\begin{tabular}{lccc}
\hline
\textbf{Tissue} & \textbf{Median (Pa)} & \textbf{IQR (Pa)} & \textbf{A-D statistic} \\
\hline
Bone                & $3.148 \times 10^{9}$ & $2.351 \times 10^{9}$ -- $3.787 \times 10^{9}$ & $0.861^{*}$ \\
Tendon              & $2.068 \times 10^{9}$ & $1.535 \times 10^{9}$ -- $2.556 \times 10^{9}$ & $0.270$ \\
Enthesis/Cartilage  & $4.930 \times 10^{8}$ & $3.795 \times 10^{8}$ -- $6.340 \times 10^{8}$ & $0.319$ \\
\hline
\end{tabular}

\vspace{1em}

\begin{tabular}{lcc}
\hline
\textbf{Comparison} & \textbf{U statistic} & \textbf{Corrected \textit{p}-value} \\
\hline
Bone vs.\ Tendon              & $311.0$ & $p = 0.008^{**}$     \\
Bone vs.\ Enthesis/Cartilage  & $399.0$ & $p < 0.0001^{****}$  \\
Tendon vs.\ Enthesis/Cartilage & $2.0$  & $p < 0.0001^{****}$  \\
\hline
\end{tabular}
\footnotesize{$^{*}$Fails Anderson-Darling normality test at $5\%$ significance 
level (critical value $= 0.721$). Wilcoxon rank-sum tests applied with Bonferroni 
correction for $n = 3$ comparisons. $^{**}p < 0.01$, $^{****}p < 0.0001$.}
\end{table}

\begin{figure} [h!]
    \includesvg[width=1\textwidth, inkscapelatex=false]{Figures/fig-aivia-enthesis.svg} 
    \caption{\textit{SHAPES improves tissue class differentiation over ROI sampling }. Comparison of ability to differentiate tissue classes within zones of interest using the (a,b)) traditional method of user selected n=20 pixel ROIs and (c,d) the SHAPES method capturing over n=10000 pixels per class. Scale bars are 10 microns. Arrows indicate location of the enthesis double tidemark}
    \label{aivia-results}
\end{figure}

Tissue segmentation performed on the AFM Young's modulus maps using the pixel classifier trained to distinguish three classes (Enthesis, Tendon, and Bone) was first evaluated by inspecting confidence maps (Fig~\ref{aivia}b) and performing iterative refinement of pixel-level class assignments until visually satisfactory separation was achieved. Final segmentation was generated using the smart segment output, with detection threshold, partition factor, and kernel size adjusted to accurately capture tissue boundaries at the scale of the modulus maps. Quantitative measurements, including per-class elasticity map (Fig~\ref{aivia}c,d,e) distributions, were extracted via the Exploration tab and compiled into easy to read summary reports.

Looking first at the traditional method of user-selected spot-based ROI AFM analysis, the Young's modulus values from each tissue type are summarized in Table~\ref{tab:wilcox_spot}. Pairwise Wilcoxon rank-sum tests with Bonferroni correction reveal all three tissue classes were significantly different from one another, confirming that SHG-guided spot-based AFM analysis resolves the known hierarchical stiffness gradient from soft fibrocartilage through to mineralized bone at the enthesis as shown in the literature,\cite{Tits2021-jk} however, the bone vs tendon comparison was less confident in comparison. Zero-valued pixels corresponding to background were 
excluded prior to analysis. Prior to comparison, data were assessed for normality using the Anderson-Darling test, but surprisingly only bone failed this criterion at the 5\% significance level and exhibited a right-skewed distribution consistent with the broad stiffness range characteristic of mineralized tissue.

In comparison, Young's modulus values extracted from the SHAPES segmented mapped ROI are summarized in Table~\ref{tab:wilcox_shapes}. Here, all three tissue classes failed the Anderson-Darling normality test, more consistent with the broad, skewed stiffness distributions expected across heterogeneous biological tissue. Pairwise Wilcoxon rank-sum tests with Bonferroni correction were therefore appropriately applied across all three class combinations. All comparisons were significant at $p<0.0001$ with the enthesis exhibiting the lowest median stiffness, followed by tendon and bone. The large pixel counts per class scale with the relative tissue area within the imaged field of view, and represent an approximately three orders-of-magnitude increase in sample size compared to the spot-based ROI approach ($n = 10^4–10^5$ vs $n = 20$ pixels per class, shown in Table~\ref{tab:wilcox_shapes} vs. Table~\ref{tab:wilcox_spot}). This dense, pixel-wise sampling improves the precision with which distributional properties are estimated and enhances sensitivity to inter-class differences, particularly in the presence of broad, non-Gaussian stiffness distributions characteristic of heterogeneous biological tissues. Consequently, the SHAPES classifier framework yields a more statistically robust and unbiased representation of tissue-level mechanical heterogeneity, enabling high-throughput AFM elasticity mapping across structurally complex interfaces such as the enthesis.

\begin{table}[h!]
\centering
\caption{Young's modulus measurements from SHAPES 
pixel classifier map and descriptive statistics from pairwise Wilcoxon rank-sum paired tests (Bonferroni corrected) . }
\label{tab:wilcox_shapes}
\begin{tabular}{lcccc}
\hline
\textbf{Tissue} & \textbf{n (pixels)} & \textbf{Median (Pa)} & \textbf{IQR (Pa)} & \textbf{A-D statistic} \\
\hline
Bone               & 29,751 & $2.591 \times 10^{9}$ & $1.447 \times 10^{9}$ -- $4.045 \times 10^{9}$ & $107.3^{*}$ \\
Tendon             & 81,687 & $1.579 \times 10^{9}$ & $1.128 \times 10^{9}$ -- $2.156 \times 10^{9}$ & $60.7^{*}$  \\
Enthesis/Cartilage & 11,379 & $6.349 \times 10^{8}$ & $3.503 \times 10^{8}$ -- $1.141 \times 10^{9}$ & $249.0^{*}$ \\
\hline
\end{tabular}

\vspace{1em}

\begin{tabular}{lcc}
\hline
\textbf{Comparison} & \textbf{U statistic} & \textbf{Corrected \textit{p}-value} \\
\hline
Bone vs.\ Tendon               & $7.531 \times 10^{8}$ & $p < 0.0001^{****}$ \\
Bone vs.\ Enthesis/Cartilage   & $5.028 \times 10^{7}$ & $p < 0.0001^{****}$ \\
Tendon vs.\ Enthesis/Cartilage & $1.861 \times 10^{8}$ & $p < 0.0001^{****}$ \\
\hline
\end{tabular}
\footnotesize{$^{*}$All three classes fail the Anderson-Darling normality 
test (critical value $= 0.752$), confirming the appropriateness of 
non-parametric testing. Wilcoxon rank-sum tests applied with Bonferroni 
correction for $n = 3$ comparisons. $^{****}p < 0.0001$.}
\end{table}

\subsection{SHAPES Generalizes to Skeletal Muscle: Endomysium and Myofiber Compartments Resolved by SHG-Guided AFM}

To demonstrate that the SHAPES workflow generalizes beyond the enthesis, the same four-step protocol was applied to a cryosectioned skeletal muscle biopsy. The collagen-rich endomysium network surrounding individual myofiber compartments within a vast array of myofiber bundles is readily visualized in the SHG channel without fixation or staining, providing an anatomically specific contrast mechanism (Fig. ~\ref{aivia-results-myo}a). A 40x region was selected from the SHG overview for AFM coregistration (Fig.~\ref{aivia-results-myo}b,c), and the resulting two-channel composite was used to train the pixel classifier with painted annotations distinguishing endomysium from myofiber compartments (Fig.~\ref{aivia-results-myo}d,e). Young's modulus maps segmented by tissue class reveal a finding which may contradict the prevailing assumption that collagen-rich regions are mechanically stiffer than surrounding parenchyma.\cite{Gillies2011-zi, Smith2011-tb} The endomysium exhibits a median stiffness of $3.975 \times 10^{6} Pa$, approximately 3-fold softer than the myofibers at $1.505 \times 10^{7} Pa$, a difference which is highly significant (Wilcoxon rank-sum, $p<0.0001$; Table \ref{tab:muscle}, Fig~\ref{aivia-results-myo}e). Both tissue classes failed the Anderson-Darling normality test, consistent with the heterogeneous stiffness distributions seen across the enthesis data and confirming the appropriateness of non-parametric testing. The segmented elasticity heatmaps make this mechanical inversion visually unambiguous: the endomysial network, bright in SHG, is consistently dark in the modulus map relative to the myofiber it surrounds (Fig~\ref{aivia-results-myo}f,g). This result, obtained using the identical instrument configuration, sample preparation, and analysis pipeline as the enthesis, demonstrates that SHAPES captures mechanobiologically meaningful structure-function relationships that would be inaccessible to structural whole myofiber bundle measurements alone. Additionally, no prior study appears to have used AFM force mapping with compartment-specific segmentation to directly compare endomysium and myofiber stiffness at the pixel level in native tissue sections, and such additional metrics may challenge assumptions that are commonly made when interpreting collagen content as a proxy for tissue stiffness in fibrotic conditions.

\begin{figure} [h!]
    \includesvg[width=1\textwidth, inkscapelatex=false]{Figures/fig-aivia-myo2.svg} 
    \caption{\textit{SHAPES applied to skeletal muscle reveals that collagen-rich endomysium is mechanically softer than the myofiber compartments it surrounds}. (a) Low-magnification SHG/2PE overview of a cryosectioned mouse skeletal muscle biopsy showing the collagen-rich endomysium network (green, SHG) surrounding individual myofiber bundles of the myofiber (magenta, 2PE autofluorescence). (b) SHG (top) and corresponding AFM (bottom) Young's modulus map of the same region. (c) Two-channel overlay composite used for pixel classifier training, showing SHG signal (green), AFM modulus map (grey), with user-annotated painted regions identifying endomysium (red) and myofiber (yellow) classes. (d) Final segmented objects output from the SHAPES pixel classifier, with endomysium (yellow-green) and myofiber (blue) classes. Violin plots (e) comparing the per pixel elasticity measurements from segmented AFM Young's modulus heatmaps for the endomysium/ECM compartment (f) and myofiber compartments (g). All scale bars represent 2 microns (inferno lookup tables, 0–50 MPa) Contrary to the prevailing assumption that increased collagen density correlates with increased stiffness, endomysium is significantly softer than the surrounding myofibers ($p < 0.0001$, Wilcoxon rank-sum test)}
    \label{aivia-results-myo}
\end{figure}

\begin{table}[h]
\caption{Young's modulus measurements from SHAPES pixel classifier segmentation of skeletal muscle and descriptive statistics from pairwise Wilcoxon rank-sum test (Bonferroni corrected).}
\label{tab:muscle}
\begin{tabular}{lllll}
\hline
\textbf{Tissue} & \textbf{n (pixels)} & \textbf{Median (Pa)} & \textbf{IQR (Pa)} & \textbf{A-D statistic} \\
\hline
Myofiber        & 30{,}210 & $1.505 \times 10^{7}$ & $8.915 \times 10^{6}$ -- $2.175 \times 10^{7}$ & $105.5^{*}$ \\
Endomysium/ECM  & 9{,}947  & $3.975 \times 10^{6}$ & $1.291 \times 10^{6}$ -- $1.256 \times 10^{7}$ & $606.5^{*}$ \\
\hline
\end{tabular}

\smallskip
\begin{tabular}{lll}
\hline
\textbf{Comparison} & \textbf{U statistic} & \textbf{Corrected \textit{p}-value} \\
\hline
Myofiber vs.\ Endomysium/ECM & $7.205 \times 10^{7}$ & $p < 0.0001^{****}$ \\
\hline
\end{tabular}

\smallskip
{\footnotesize $^{*}$Both classes fail the Anderson-Darling normality test (critical value $= 0.752$), confirming the appropriateness of non-parametric testing with Wilcoxon rank-sum test.}
\end{table}


\section{Discussion}\label{sec4}

When directly comparing the results of the large-kernel pixel classifier (Fig~\ref{aivia-results}(c,d)) to the traditional method of user-selected spot-based ROIs (Fig~\ref{aivia-results}(a,b)), both approaches recover the expected mechanical hierarchy, with enthesis/cartilage exhibiting lower stiffness than the surrounding tendon and bone regions, consistent with the known biomechanical organization of the fibrocartilaginous enthesis \cite{Benjamin2002-iz, Tits2021-jk}. Interestingly, the Anderson-Darling normality tests pass for the enthesis and tendon classes in the user selected spot-based ROI, but fail for the segmented SHAPES maps, highlighting how the reliance on manually selected spot-based ROI introduces the risk of confirmation bias through selective sampling. Similarly, the bone vs tendon Bonferroni-corrected p-value was slightly less significant than the fully mapped SHAPES data, further displaying how the small n=20 number of pixel counts obtained by this approach may be insufficient to satisfy assumptions of normality when comparing smaller expected changes between experimental cohorts, potentially limiting the robustness of downstream parametric statistical inference. \cite{Stylianou2019-al, Cho2023-rt}

In contrast, the SHAPES pixel classifier enables extraction of an elasticity metric from AFM and a structural metric from SHG for every measured pixel within a full mapped field of view, yielding substantially larger per-class sample sizes and producing distributions which more realistically represent the mechanical heterogeneity of each tissue compartment \cite{Liu2020-uo, Khattignavong2023-xs}. This approach prevents biased ROI selection and increases statistical power, as reflected in the Wilcoxon rank-sum test results summarized in Tables~\ref{tab:wilcox_spot} and~\ref{tab:wilcox_shapes}. Notably, all pairwise comparisons remain significant at p$<$0.0001 in the SHAPES analysis, and the separation between tissue classes is consistent with prior correlative structural-mechanical characterization of collagenous tissues using complementary imaging modalities \cite{Plekhanov2023-yb, Arshee2025-wg, Cisek2021-fy}. While not explored in the present work, this framework also enables further intra-class segmentation to investigate phenotypic heterogeneity. For example, collagen alignment metrics derived from the SHG channel could be correlated with local mechanical properties by segmenting individual fibers, as demonstrated in prior work.\cite{Shi2024-az} Additionally, the enthesis region could be further subdivided to assess spatial differences between the bone-facing and tendon-facing edges of the interface, particularly across the enthesis zone, observed as a double tidemark\cite{Camy2022-gs} in this image, and used to correlate directly to AFM and potentially other modalities such as fluorescence lifetime imaging\cite{Jeong2018-rf} or sparse spectral stimulated Raman imaging\cite{Pence2021-zo}, or better yet, these plus AFM all on the same sample.\cite{Quansah2023-um} These type of insights greatly inform the design of engineered 3D-printable biocompatible substrates as well by providing both structural and mechanical information from the same imaging pixel.

The applicability of this workflow to other clinical applications is directly demonstrated in Fig.~\ref{aivia-results-myo}, where SHAPES was applied without modification to a cryosectioned skeletal muscle biopsy,  revealing that the collagen-rich endomysium network is approximately 3-fold softer than the myofibers (median $3.975 \times 10^6 Pa$ vs. $1.505 \times 10^7 Pa, p < 0.0001$; Table 3), a finding that directly contradicts the assumption that collagen-dense tissue compartments are mechanically stiffer than the myofiber within. This is consistent with prior work demonstrating that passive mechanical stiffness in typically developing and cerebral palsy muscles are governed by ECM-level changes that cannot be inferred from full fiber-level measurements alone\cite{Smith2011-tb} and provides the first pixel-resolved, label-free spatial confirmation of this compartment-level mechanical inversion in native tissue.

\section{Conclusion \& Outlook}\label{sec5}

The label-free correlative imaging workflow described in this work combines SHG and 2PE microscopy with atomic force microscopy (AFM) to assess collagen microstructure and local stiffness in native, unfixed tissue. By combining extended range AFM with SHG-guided ROI selection, SHAPES substantially reduces the operator burden associated with navigating complex tissue topographies, minimizes the risk of broken AFM probes or stopped scans, and makes AFM analysis of heterogeneous tissue samples practically achievable for core facilities and clinical laboratories. AI-based pixel classification correlated across SHG and AFM datasets further enhances the analytical throughput and objectivity of the workflow, eliminating the confirmation bias inherent to manual ROI selection and enabling statistically robust, macroscale mechanical phenotyping. SHAPES is also fully compatible with downstream fixation and staining, allowing colocalization with established histological and molecular markers to contextualize mechanical findings within existing literature.

This is of particular significance in pathological contexts where the relationship between collagen content and tissue mechanics is assumed but rarely directly tested. While fibrosis of the enthesis (the site where tendons and ligaments attach to bone) represents just one manifestation of collagen-driven pathology, fibrosis in general is characterized by the overproduction and disorganization of collagen across a wide range of tissue types and provides a compelling example of where SHAPES can add diagnostic value. The prevailing assumption that increased collagen density corresponds to increased tissue stiffness can now be directly interrogated at the microstructural level rather than inferred from structural data alone. Indeed, ongoing work applying SHAPES to muscle fibrosis in myofiber bundles from muscle biopsies has revealed that large fibrotic formations, traditionally expected to stiffen tissue, are in fact significantly softer than the surrounding parenchyma, a finding which challenges established hypotheses about the mechanical consequences of excessive fibrosis and opens new lines of inquiry into its pathophysiology. The ability of SHAPES to generate this class of structure-function data across diverse tissue types, demonstrated here using a single unified protocol, positions it as a broadly applicable platform for mechanobiological hypothesis testing in both preclinical and translational settings. Overall, SHAPES improves anatomical specificity, reproducibility, and provides a user-friendly workflow for nanomechanical phenotyping of collagen-rich extracellular matrices in preclinical and translational contexts, without the need for exogenous labels, specialized sample preparation, or custom software. 

\backmatter

\bmhead{Supplementary information}

Root mean square error distribution plots for AFM plots can be found in supplemental Figure \ref{suppRMS}. All raw images, datasets, pixel classifier and training sets can be found on the ALMS Github repository.

\bmhead{Acknowledgements}

All instruments used for this work are available for public use at the Advanced Light Microscopy/Spectroscopy Laboratory and Leica Microsystems Center of Excellence at the California NanoSystems Institute at UCLA (RRID:SCR 022789) and the Nano and Pico Characterization Laboratory (RRID:SCR 022924) housed within the California NanoSystems Institute (CNSI) at the University of California, Los Angeles, with funding support from NIH Shared Instrumentation Grant S10OD025017.

\section*{Declarations}

The authors have no conflicts of interest to declare. 

\begin{appendices}

\section{Supplementary Information}\label{secA1}

\textbf{Supplemental Video 1.} SHG/2PE image alignment within the AFM control software. The SHG/2PE maximum projection is imported directly into the JPK NanoWizard 4A control software with physical dimensions entered via the manual scaling feature to match the AFM coordinate system. The image is rotated 90\degree and overlaid at 50\% transparency onto the live brightfield image acquired through the integrated Leica M205 FCA stereomicroscope using the Optical Transform function in the JPK SPM Desktop Viewer Window. Precise coregistration is achieved by aligning shared structural landmarks visible in both modalities, including tissue edges and distinct morphological features of the sample. The completed overlay is then used to identify and select the target region of interest for subsequent AFM force mapping.

\begin{figure}[h]
    \includesvg[width=0.5\textwidth, inkscapelatex=false]{Figures/rms.svg} 
    \caption{Supplementary Figure 1: RMS distribution for all AFM curves in map. Root mean square is a useful metric for evaluating accuracy of fit for AFM force curves. Across all pixels in the map, the median residual error of 18nN is negligible.}
    \label{suppRMS}
\end{figure}

\newpage
%%=============================================%%
%% For submissions to Nature Portfolio Journals %%
%% please use the heading ``Extended Data''.   %%
%%=============================================%%

%%=============================================================%%
%% Sample for another appendix section			       %%
%%=============================================================%%

%% \section{Example of another appendix section}\label{secA2}%
%% Appendices may be used for helpful, supporting or essential material that would otherwise 
%% clutter, break up or be distracting to the text. Appendices can consist of sections, figures, 
%% tables and equations etc.

\end{appendices}

%%===========================================================================================%%
%% If you are submitting to one of the Nature Portfolio journals, using the eJP submission   %%
%% system, please include the references within the manuscript file itself. You may do this  %%
%% by copying the reference list from your .bbl file, paste it into the main manuscript .tex %%
%% file, and delete the associated \verb+\bibliography+ commands.                            %%
%%===========================================================================================%%

\bibliography{refs/paperpile}% common bib file
%% if required, the content of .bbl file can be included here once bbl is generated
%%\input sn-article.bbl

\end{document}
