## Supplementary material for "Correlative SHG-AFM imaging workflow for label-free quantitative analysis of collagen structure–function relationships": LaTEX: springer nature template.pdf

### Article Title

First Author[1,2*], Second Author[2,3†] and Third Author[1,2†]

[1*]Department, Organization, Street, City, 100190, State, Country.
[2]Department, Organization, Street, City, 10587, State, Country.
[3]Department, Organization, Street, City, 610101, State, Country.

Contributing authors:;
†These authors contributed equally to this work.

**Abstract**

The abstract serves both as a general introduction to the topic and as a brief, non-technical summary of the main results and their implications. Authors are advised to check the author instructions for the journal they are submitting to for word limits and if structural elements like subheadings, citations, or equations are permitted.

**Keywords:** keyword1, Keyword2, Keyword3, Keyword4

#### 1 Introduction

The Introduction section, of referenced text Campbell and Gear (1995) expands on the background of the work (some overlap with the Abstract is acceptable). The introduction should not include subheadings.

Springer Nature does not impose a strict layout as standard however authors are advised to check the individual requirements for the journal they are planning to submit to as there may be journal-level preferences. When preparing your text please also be aware that some stylistic choices are not supported in full text XML (publication version), including coloured font. These will not be replicated in the typeset article if it is accepted.

#### 2 Results

Sample body text. Sample body text. Sample body text. Sample body text. Sample body text. Sample body text. Sample body text. Sample body text.

#### 3 This is an example for first level head—section head

##### 3.1 This is an example for second level head—subsection head

###### 3.1.1 This is an example for third level head---subsubsection head

Sample body text. Sample body text. Sample body text. Sample body text. Sample body text. Sample body text. Sample body text. Sample body text.

#### 4 Equations

Equations in LaTeX can either be inline or on-a-line by itself ("display equations"). For inline equations use the `$...$` commands. E.g.: The equation $H\psi = E\psi$ is written via the command `$H \psi = E \psi$`.

For display equations (with auto generated equation numbers) one can use the equation or align environments:

$$\|\tilde{X}(k)\|^2 \leq \frac{\sum\limits_{i=1}^{p} \left\|\tilde{Y}_i(k)\right\|^2 + \sum\limits_{j=1}^{q} \left\|\tilde{Z}_j(k)\right\|^2}{p+q}. \tag{1}$$

where,

$$\begin{aligned} D_\mu &= \partial_\mu - ig\frac{\lambda^a}{2}A^a_\mu \\ F^a_{\mu\nu} &= \partial_\mu A^a_\nu - \partial_\nu A^a_\mu + g f^{abc} A^b_\mu A^a_\nu \end{aligned} \tag{2}$$

Notice the use of `\nonumber` in the align environment at the end of each line, except the last, so as not to produce equation numbers on lines where no equation numbers are required. The `\label{}` command should only be used at the last line of an align environment where `\nonumber` is not used.

$$Y_\infty = \left(\frac{m}{\text{GeV}}\right)^{-3} \left[1 + \frac{3\ln(m/\text{GeV})}{15} + \frac{\ln(c_2/5)}{15}\right] \tag{3}$$

The class file also supports the use of `\mathbb{}`, `\mathscr{}` and `\mathcal{}` commands. As such `\mathbb{R}`, `\mathscr{R}` and `\mathcal{R}` produces $\mathbb{R}$, $\mathscr{R}$ and $\mathcal{R}$ respectively (refer Subsubsection 3.1.1).

### 5 Tables

Tables can be inserted via the normal table and tabular environment. To put footnotes inside tables you should use `\footnotetext[]{...}` tag. The footnote appears just below the table itself (refer Tables 1 and 2). For the corresponding footnotemark use `\footnotemark[...]`

**Table 1** Caption text

| Column 1 | Column 2 | Column 3 | Column 4 |
|---|---|---|---|
| row 1 | data 1 | data 2 | data 3 |
| row 2 | data 4 | data 5[1] | data 6 |
| row 3 | data 7 | data 8 | data 9[2] |

Source: This is an example of table footnote. This is an example of table footnote.

[1]Example for a first table footnote. This is an example of table footnote.

[2]Example for a second table footnote. This is an example of table footnote.

The input format for the above table is as follows:

```
\begin{table}[<placement-specifier>]
\caption{<table-caption>}\label{<table-label>}%
\begin{tabular}{@{}llll@{}}
\toprule
Column 1 & Column 2 & Column 3 & Column 4\\
\midrule
row 1 & data 1 & data 2  & data 3 \\
row 2 & data 4 & data 5\footnotemark[1] & data 6 \\
row 3 & data 7 & data 8  & data 9\footnotemark[2]\\
\botrule
\end{tabular}
\footnotetext{Source: This is an example of table footnote.
This is an example of table footnote.}
\footnotetext[1]{Example for a first table footnote.
This is an example of table footnote.}
\footnotetext[2]{Example for a second table footnote.
This is an example of table footnote.}
\end{table}
```

In case of double column layout, tables which do not fit in single column width should be set to full text width. For this, you need to use `\begin{table*} ...`

**Table 2** Example of a lengthy table which is set to full textwidth

| | Element 1[1] | | | Element 2[2] | | |
|---|---|---|---|---|---|---|
| Project | Energy | $\sigma_{calc}$ | $\sigma_{expt}$ | Energy | $\sigma_{calc}$ | $\sigma_{expt}$ |
| Element 3 | 990 A | 1168 | $1547 \pm 12$ | 780 A | 1166 | $1239 \pm 100$ |
| Element 4 | 500 A | 961 | $922 \pm 10$ | 900 A | 1268 | $1092 \pm 40$ |

Note: This is an example of table footnote. This is an example of table footnote this is an example of table footnote this is an example of table footnote this is an example of table footnote.

[1]Example for a first table footnote.

[2]Example for a second table footnote.

`\end{table*}` instead of `\begin{table} ... \end{table}` environment. Lengthy tables which do not fit in textwidth should be set as rotated table. For this, you need to use `\begin{sidewaystable} ... \end{sidewaystable}` instead of `\begin{table*} ... \end{table*}` environment. This environment puts tables rotated to single column width. For tables rotated to double column width, use `\begin{sidewaystable*} ... \end{sidewaystable*}`.

### 6 Figures

As per the LaTeX standards you need to use eps images for LaTeX compilation and `pdf/jpg/png` images for `PDFLaTeX` compilation. This is one of the major difference between LaTeX and `PDFLaTeX`. Each image should be from a single input .eps/vector image file. Avoid using subfigures. The command for inserting images for LaTeX and `PDFLaTeX` can be generalized. The package used to insert images in `LaTeX/PDFLaTeX` is the graphicx package. Figures can be inserted via the normal figure environment as shown in the below example:

```
\begin{figure}[<placement-specifier>]
\centering
\includegraphics{<eps-file>}
\caption{<figure-caption>}\label{<figure-label>}
\end{figure}
```

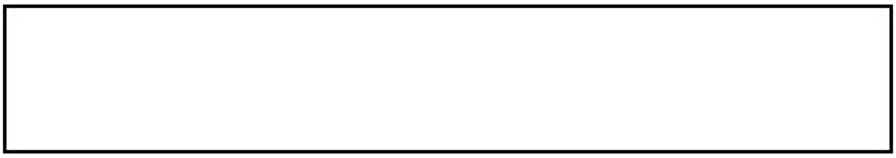

**Fig. 1** This is a widefig. This is an example of long caption this is an example of long caption this is an example of long caption this is an example of long caption

**Table 3** Tables which are too long to fit, should be written using the "sidewaystable" environment as shown here

| | Element 1[1] | | | Element[2] | | |
|---|---|---|---|---|---|---|
| Projectile | Energy | $\sigma_{calc}$ | $\sigma_{expt}$ | Energy | $\sigma_{calc}$ | $\sigma_{expt}$ |
| Element 3 | 990 A | 1168 | $1547 \pm 12$ | 780 A | 1166 | $1239 \pm 100$ |
| Element 4 | 500 A | 961 | $922 \pm 10$ | 900 A | 1268 | $1092 \pm 40$ |
| Element 5 | 990 A | 1168 | $1547 \pm 12$ | 780 A | 1166 | $1239 \pm 100$ |
| Element 6 | 500 A | 961 | $922 \pm 10$ | 900 A | 1268 | $1092 \pm 40$ |

Note: This is an example of table footnote this is an example of table footnote this is an example of table footnote this is an example of table footnote this is an example of table footnote.

[1] This is an example of table footnote.

In case of double column layout, the above format puts figure captions/images to single column width. To get spanned images, we need to provide `\begin{figure*}` ... `\end{figure*}`.

For sample purpose, we have included the width of images in the optional argument of `\includegraphics` tag. Please ignore this.

### 7 Algorithms, Program codes and Listings

Packages `algorithm`, `algorithmicx` and `algpseudocode` are used for setting algorithms in LaTeX using the format:

```
\begin{algorithm}
\caption{<alg-caption>}\label{<alg-label>}
\begin{algorithmic}[1]
. . .
\end{algorithmic}
\end{algorithm}
```

You may refer above listed package documentations for more details before setting `algorithm` environment. For program codes, the "verbatim" package is required and the command to be used is `\begin{verbatim}` ... `\end{verbatim}`.

Similarly, for `listings`, use the `listings` package. `\begin{lstlisting}` ... `\end{lstlisting}` is used to set environments similar to `verbatim` environment. Refer to the `lstlisting` package documentation for more details.

A fast exponentiation procedure:

```
begin
  for i := 1 to 10 step 1 do
      expt(2, i);
      newline() od                  Comments will be set flush to the right margin
where
proc expt(x, n) ≡
  z := 1;
  do if n = 0 then exit fi;
     do if odd(n) then exit fi;
        comment: This is a comment statement;
        n := n/2; x := x * x od;
     { n > 0 };
     n := n - 1; z := z * x od;
  print(z).
end
```

**Algorithm 1** Calculate $y = x^n$

**Require:** $n \geq 0 \vee x \neq 0$
**Ensure:** $y = x^n$
1: $y \Leftarrow 1$
2: **if** $n < 0$ **then**
3: $\quad X \Leftarrow 1/x$
4: $\quad N \Leftarrow -n$
5: **else**
6: $\quad X \Leftarrow x$
7: $\quad N \Leftarrow n$
8: **end if**
9: **while** $N \neq 0$ **do**
10: $\quad$ **if** $N$ is even **then**
11: $\qquad X \Leftarrow X \times X$
12: $\qquad N \Leftarrow N/2$
13: $\quad$ **else**[$N$ is odd]
14: $\qquad y \Leftarrow y \times X$
15: $\qquad N \Leftarrow N - 1$
16: $\quad$ **end if**
17: **end while**

```
for i:=maxint to 0 do
begin
{ do nothing }
end;
Write('Case-insensitive-');
Write('Pascal-keywords.');
```

### 8 Cross referencing

Environments such as figure, table, equation and align can have a label declared via the `\label{#label}` command. For figures and table environments use the `\label{}` command inside or just below the `\caption{}` command. You can then use the `\ref{#label}` command to cross-reference them. As an example, consider the label declared for Figure 1 which is `\label{fig1}`. To cross-reference it, use the command `Figure \ref{fig1}`, for which it comes up as "Figure 1".

To reference line numbers in an algorithm, consider the label declared for the line number 2 of Algorithm 1 is `\label{algln2}`. To cross-reference it, use the command `\ref{algln2}` for which it comes up as line 2 of Algorithm 1.

#### 8.1 Details on reference citations

Standard LaTeX permits only numerical citations. To support both numerical and author-year citations this template uses `natbib` LaTeX package. For style guidance please refer to the template user manual.

Here is an example for `\cite{...}`: Campbell and Gear (1995). Another example for `\citep{...}`: (Slifka and Whitton 2000). For author-year citation mode, `\cite{...}` prints Jones et al. (1990) and `\citep{...}` prints (Jones et al., 1990).

All cited bib entries are printed at the end of this article: Hamburger (1995), Geddes et al. (1992), Broy (1992), Seymour (1981), Smith (1976), Chung and Morris (1978), Hao et al. (2014), Babichev et al. (2002), Beneke et al. (1997), Stahl (2020) and Abbott et al. (2019).

### 9 Examples for theorem like environments

For theorem like environments, we require `amsthm` package. There are three types of predefined theorem styles exists—`thmstyleone`, `thmstyletwo` and `thmstylethree`

| | |
|---|---|
| `thmstyleone` | Numbered, theorem head in bold font and theorem text in italic style |
| `thmstyletwo` | Numbered, theorem head in roman font and theorem text in italic style |
| `thmstylethree` | Numbered, theorem head in bold font and theorem text in roman style |

For mathematics journals, theorem styles can be included as shown in the following examples:

**Theorem 1** (Theorem subhead) *Example theorem text. Example theorem text. Example theorem text. Example theorem text. Example theorem text. Example theorem text. Example theorem text. Example theorem text. Example theorem text. Example theorem text. Example theorem text.*

Sample body text. Sample body text. Sample body text. Sample body text. Sample body text. Sample body text. Sample body text. Sample body text.

**Proposition 2** *Example proposition text. Example proposition text. Example proposition text. Example proposition text. Example proposition text. Example proposition text. Example proposition text. Example proposition text. Example proposition text. Example proposition text.*

Sample body text. Sample body text. Sample body text. Sample body text. Sample body text. Sample body text. Sample body text. Sample body text.

*Example 1* Phasellus adipiscing semper elit. Proin fermentum massa ac quam. Sed diam turpis, molestie vitae, placerat a, molestie nec, leo. Maecenas lacinia. Nam ipsum ligula, eleifend at, accumsan nec, suscipit a, ipsum. Morbi blandit ligula feugiat magna. Nunc eleifend consequat lorem.

Sample body text. Sample body text. Sample body text. Sample body text. Sample body text. Sample body text. Sample body text. Sample body text.

*Remark 1* Phasellus adipiscing semper elit. Proin fermentum massa ac quam. Sed diam turpis, molestie vitae, placerat a, molestie nec, leo. Maecenas lacinia. Nam ipsum ligula, eleifend at, accumsan nec, suscipit a, ipsum. Morbi blandit ligula feugiat magna. Nunc eleifend consequat lorem.

Sample body text. Sample body text. Sample body text. Sample body text. Sample body text. Sample body text. Sample body text. Sample body text.

**Definition 1** (Definition sub head) Example definition text. Example definition text. Example definition text. Example definition text. Example definition text. Example definition text. Example definition text. Example definition text.

Additionally a predefined "proof" environment is available: `\begin{proof} ... \end{proof}`. This prints a "Proof" head in italic font style and the "body text" in roman font style with an open square at the end of each proof environment.

*Proof* Example for proof text. Example for proof text. Example for proof text. Example for proof text. Example for proof text. Example for proof text. Example for proof text. Example for proof text. Example for proof text. Example for proof text. □

Sample body text. Sample body text. Sample body text. Sample body text. Sample body text. Sample body text. Sample body text. Sample body text.

*Proof of Theorem 1* Example for proof text. Example for proof text. Example for proof text. Example for proof text. Example for proof text. Example for proof text. Example for proof text. Example for proof text. Example for proof text. Example for proof text. □

For a quote environment, use `\begin{quote}...\end{quote}`

> Quoted text example. Aliquam porttitor quam a lacus. Praesent vel arcu ut tortor cursus volutpat. In vitae pede quis diam bibendum placerat. Fusce elementum convallis neque. Sed dolor orci, scelerisque ac, dapibus nec, ultricies ut, mi. Duis nec dui quis leo sagittis commodo.

Sample body text. Sample body text. Sample body text. Sample body text. Sample body text (refer Figure 1). Sample body text. Sample body text. Sample body text (refer Table 3).

#### 10 Methods

Topical subheadings are allowed. Authors must ensure that their Methods section includes adequate experimental and characterization data necessary for others in the field to reproduce their work. Authors are encouraged to include RIIDs where appropriate.

**Ethical approval declarations** (only required where applicable) Any article reporting experiment/s carried out on (i) live vertebrate (or higher invertebrates), (ii) humans or (iii) human samples must include an unambiguous statement within the methods section that meets the following requirements:

1. Approval: a statement which confirms that all experimental protocols were approved by a named institutional and/or licensing committee. Please identify the approving body in the methods section
2. Accordance: a statement explicitly saying that the methods were carried out in accordance with the relevant guidelines and regulations
3. Informed consent (for experiments involving humans or human tissue samples): include a statement confirming that informed consent was obtained from all participants and/or their legal guardian/s

If your manuscript includes potentially identifying patient/participant information, or if it describes human transplantation research, or if it reports results of a clinical trial then additional information will be required. Please visit (https://www.nature.com/nature-research/editorial-policies) for Nature Portfolio journals, (https://www.springer.com/gp/authors-editors/journal-author/journal-author-helpdesk/publishing-ethics/14214) for Springer Nature journals, or (https://www.biomedcentral.com/getpublished/editorial-policies#ethics+and+consent) for BMC.

#### 11 Discussion

Discussions should be brief and focused. In some disciplines use of Discussion or 'Conclusion' is interchangeable. It is not mandatory to use both. Some journals prefer a section 'Results and Discussion' followed by a section 'Conclusion'. Please refer to Journal-level guidance for any specific requirements.

#### 12 Conclusion

Conclusions may be used to restate your hypothesis or research question, restate your major findings, explain the relevance and the added value of your work, highlight any limitations of your study, describe future directions for research and recommendations.

In some disciplines use of Discussion or 'Conclusion' is interchangeable. It is not mandatory to use both. Please refer to Journal-level guidance for any specific requirements.

**Supplementary information.** If your article has accompanying supplementary file/s please state so here.

Authors reporting data from electrophoretic gels and blots should supply the full unprocessed scans for key as part of their Supplementary information. This may be requested by the editorial team/s if it is missing.

Please refer to Journal-level guidance for any specific requirements.

**Acknowledgements.** Acknowledgements are not compulsory. Where included they should be brief. Grant or contribution numbers may be acknowledged.

Please refer to Journal-level guidance for any specific requirements.

### Declarations

Some journals require declarations to be submitted in a standardised format. Please check the Instructions for Authors of the journal to which you are submitting to see if you need to complete this section. If yes, your manuscript must contain the following sections under the heading 'Declarations':

- Funding
- Conflict of interest/Competing interests (check journal-specific guidelines for which heading to use)
- Ethics approval and consent to participate
- Consent for publication
- Data availability
- Materials availability
- Code availability
- Author contribution

If any of the sections are not relevant to your manuscript, please include the heading and write 'Not applicable' for that section.

Editorial Policies for:

Springer journals and proceedings: https://www.springer.com/gp/editorial-policies

Nature Portfolio journals: https://www.nature.com/nature-research/editorial-policies

*Scientific Reports*: https://www.nature.com/srep/journal-policies/editorial-policies

BMC journals: https://www.biomedcentral.com/getpublished/editorial-policies

### Appendix A Section title of first appendix

An appendix contains supplementary information that is not an essential part of the text itself but which may be helpful in providing a more comprehensive understanding of the research problem or it is information that is too cumbersome to be included in the body of the paper.
