## Supplementary material for "Correlative SHG-AFM imaging workflow for label-free quantitative analysis of collagen structure–function relationships": LaTEX: user-manual.pdf

Springer Nature

### User Manual

#### Journal article LaTeX authoring template

### User Manual

#### Journal article LaTeX authoring template

Copyright Springer Nature
Prepared by Straive TeX Support

#### 1. Introduction

Springer Nature has developed an authoring template to help you prepare journal articles in LaTeX that conform to Springer Nature technical requirements. The LaTeX authoring template is inclusive of all research disciplines and provides key placeholders for policy requirements. Due to the universal nature of the template you are expected to read the submission guidelines and policies of the journal you are preparing your submission for and should defer to journal-level instructions.

The template is stylistically neutral. We recommend avoiding the introduction of any unnecessary formatting as non-standard packages and macros are frequent causes of error. Wherever possible please do not add further packages to the template. Adhering to these guidelines will ease the production process of any accepted manuscript and help avoid misinterpretation of your LaTeX code.

This documentation is not intended to give an introduction to LaTeX. For questions concerning TeX systems/installations or the LaTeX mark-up language in general, please visit http://tug.ctan.org/ or any other TeX user group worldwide. The essential reference for LaTeX is Mittelbach F., Goossens M. *(2004) The LaTeX Companion. 2nd edn.*, but there are many other good books about LaTeX.

If you plan to publish a book with Springer Nature please make sure to use the TeX template for books available at https://www.springernature.com/gp/authors/publish-a-book/manuscript-guidelines

#### 2. How to prepare your article

Ensure that you have LaTeX2e version installed on your computer. You are provided with a class file “sn-jnl.cls” and must keep this class file in the same directory as your manuscript files. Note that the class file depends on the following standard packages which are available along with recent LaTeX installations:

| | | | |
|---|---|---|---|
| graphicx | multirow | amsmath | amssymb |
| amsfonts | mathrsfs | rotating | appendix |
| textcomp | manyfoot | booktabs | etoolbox |
| algorithm | xcolor | algorithmicx | algpseudocode |
| listings | url | | |

The following additional packages are used in the class file for providing add-on functionalities to the template:

| | | | |
|---|---|---|---|
| geometry | apacite | hypcap | hyperref |
| breakurl | natbib | wrapfig | amsthm |

To learn more about the underlying packages, please read the respective documentations (try, e.g., texdoc [package name] at your shell prompt or visit http://tug.ctan.org/).

##### *2.1 Options*

We have supplied a LaTeX template file—"sn-article.tex" for single column layout. Should you need to switch to double column layout to suit journal-level submission requirements provide the `[iicol]` option to `\documentclass[iicol]{...}` in the preamble.

For the peer review and editorial stages you are requested to enable double-line spacing by using the `[referee]` option.

`\documentclass{sn-jnl}` – single line spaced

`\documentclass[referee]{sn-jnl}` – double line spaced

To enable line numbers in the margin, use the [lineno] option.

`\documentclass[lineno]{sn-jnl}` – prints line numbers in the margin

The sample file contains the lines for calling class files, preamble area, and the start/end of document where major elements required for an article are placed. Explanatory comments are included for each element. You can add your actual manuscript content in place of these sample elements. The standard structure of each element for an article is explained in detail in the following sections.

To use the Springer Nature authoring template put all the package files in your working directory, edit the file "sn-article.tex" in your preferred text editor, and run LaTeX as usual.

The resulting layout is optimized for peer review and will not be identical to the layout of a final article. You are not responsible for any final page layout and the use of fine-tuning commands like `\break`, `\pagebreak`, `\vspace`, `\bigskip`, `\clearpage`, *etc*. are not encouraged. Please use semantic mark-up as far as possible and avoid additional formatting commands.

##### *2.2 How to prepare your article for Nature Portfolio Journals*

Particular requirements exist for articles prepared for submission to *Nature* and the Nature Portfolio journals. Authors must check the journal-specific formatting instructions before submission and note in particular the following:

- Footnotes should not be used in *Nature*-journal submissions
- Do not upload a bibliography file. Instead include the references within the manuscript file itself. You may do this by copying the reference list from your .bbl file, and pasting it into the bibliography environment of the main manuscript .tex file
- Online only Methods should be presented in a separate section after the end of the main text and reference list
- Figure legends should be presented at the end of the file
- The reference list for extended data should be separate and carry on from the main referencing numbering

#### 3. Package features and some important settings

##### *3.1. Language*

English is the default language used for typesetting rules.

##### *3.2. Fonts*

Please refrain from using custom fonts.

*Text fonts:* Unlike the final published version, the authoring template uses non-commercial fonts: Computer Modern. These fonts are free version of the PostScript standard fonts and are supplied as part of all standard TeX distributions.

*Math fonts:* The standard Computer Modern math fonts are used.

### 4. Preamble

The preamble part comes between the document class line — `\documentclass{...}` —and the beginning of your document —`\begin{document}`. Use this preamble area to include additional packages.

### 5. Major structures/elements

Article contents are divided into three main elements—front matter, main matter, and back matter. The elements before `\maketitle` tag are considered as front matter elements and the elements placed below `\maketitle` tag are main matter elements. Back matter elements follow the \backmatter tag..

| **Front matter** | **Main matter** | **Back matter** |
|---|---|---|
| \title{. . .} | \section{. . .} | \bmhead{. . .} |
| \subtitle{. . .} | \subsection{. . .} | |
| \author{. . .} | \subsubsection{. . .} | \begin{appendices}. . .<br>\end{appendices} |
| \affil{. . .} | \begin{verbatim}. .\end{verbatim} | \bibliography{. . .} |
| \abstract{. . .} | \begin{algorithm}<br>. . .<br>\end{algorithm} | |
| \keywords{. . .} | \begin{table}. . .\end{table} | |
| \pw{. . .} | \begin{figure}. . .\end{figure} | |
| \maketitle | \begin{enumerate}<br>\end{enumerate} | |
| | \begin{unenumerate}. . .<br>\end{unenumerate} | |
| | \begin{itemize}. . .\end{itemize} | |
| | \begin{theorem}. . .\end{theorem} | |
| | \begin{definition}<br>\end{definition} | |
| | \begin{proof}. . .\end{proof} | |
| | \begin{equation}. .\end{equation} | |
| | \begin{align}...\end{align} | |

The above listed tags are pre-defined in the class file "sn-jnl.cls". If necessary, you can define your own customized tags as per your preference.

### 6. Front matter elements

The tagging details of article opener elements are as follows:

1. \title[<short-form-of-article-title>]{<article-title>}

   This tag contains two parameters. The first one is optional but {<article-title>} is mandatory.

2. \author*[<affil-num>]{<author-name>}— to be used for the corresponding author who is nominated as being responsible for the manuscript as it moves through the entire publication process.

   \author[<affil-num>]{<author-name>} —to be used for all other authors.

3. \affil[<sequence>]{<address-details>}—affiliation/address details are provided inside this tag. In case of multiple addresses, just provide sequential Arabic numerals in the optional argument of this tag. This number is used to denote the affiliation for the respective authors. In case of single author/address, this optional argument can be ignored. For example:

   \author{...}
   \affil{...}

4. \affil*[<sequence>]{<address-details>}—This tag is to represent author's present/corresponding address.

5. Other common front matter tags include:

| | | | |
|---|---|---|---|
| \email{...} | \nomail | \orgdiv{...} | \orgname{...} |
| \street{...} | \city{...} | \state{...} | \country{...} |
| \orgaddress{...} | \postcode{...} | \equalcont{...} | \abstract{...} |
| \keywords{...} | \received{...} | | |

\nomail—Provide this tag for the author without email id.

\pacs—One of keyword group.

\maketitle—This tag is mandatory to print the front matter elements in the output.

### 7. Main matter elements

##### *7.1. Section headings*

The template allows three levels of headings in different styles:

`\section{<First level heading>}` –> Numbered and printed in bold font style

`\subsection{<Second level heading>}` –> Numbered and printed in smaller bold font style

`\subsubsection{<Third level heading>}` –> Numbered and printed with further smaller bold font style

To use unnumbered level heads, provide the `\unnumbered` command in the preamble area.

##### *7.2. Mathematical formulae*

The "amsmath" package provides various features for displayed equations and other mathematical constructs and you are strongly encouraged to use the mark-ups provided by this package. Do not use manual skips to align an equation.

##### *7.3. Figures and Tables*

The standard interface for graphic inclusion is the `\includegraphics` command provided by the graphicx package. **Do not place your image files in subfolders, as the manuscript submission system will be unable to find them.** Include them on the same level as your LaTeX document. Also make sure that each figure is from a single input image file. Avoid using subfigures. LaTeX compiler accepts only .eps image file format. If you have .ps image file format, rename them to .eps images.

The format used for numbered "figures" and "tables" is similar to LaTeX basic format:

```
\begin{figure}[t]
\centering
\includegraphics{<image-file-name>}
\caption{<Caption text>}\label{...}
\end{figure}
```

In case of double column layout, the above format puts figure captions/images to single column width. To get spanned images, you need to provide
`\begin{figure*}...\end{figure*}`.

For sample purpose, we have included the width of images in the optional argument of `\includegraphics` tag. Please ignore this.

The format for a table is as follows:

```
\begin{table}[<float-position>]
\caption{...}\label{<table-label>}
\begin{tabular}{<column-alignment-preamble>}
```

```
\toprule
...  & ... & ... & ... \\
\midrule
... & ... & ... & ...\footnotemark[1] \\
... & ... & ... & ... \\
... & ... & ... & ...\footnotemark[2] \\
\botrule
\end{tabular}
\footnotetext{...}
\footnotetext[1]{...}
\end{table}
```

Command to be used for rotated tables:

`\begin{sidewaystable}...\end{sidewaystable}`

To span tables across columns in double column layout:

`\begin{table*}...\end{table*}`

To span rotated tables across columns in double column layout:

`\begin{sidewaystable*}...\end{sidewaystable*}`

To refer table footnote number in table body, use the

`\footnotemark[...]` command.

##### *7.4. Lists*

The default list commands available in LaTeX can be used to set different types of lists:

1. **numbered:** `\begin{enumerate}...\end{enumerate}`

   first level—Arabic numerals;

   second level—lowercase alphabet;

   third level—lowercase roman numerals;

2. **unnumbered:** `\begin{unenumerate}...\end{unenumerate}`

3. **custom list:** `\begin{itemize}...\end{itemize}`

   First level—bulleted; second level—dash list;

   Nested lists are allowed for numbered and custom lists .

##### *7.5. Theorem-like environments*

For theorem-like environments, we require the "amsthm" package. There are three types of predefined theorem styles exist—`thmstyleone`, `thmstyletwo`, and `thmstylethree`. The below table shows the output details for each style:

| | |
|---|---|
| thmstyleone | Numbered, theorem head in bold font and theorem text in italic style |
| thmstyletwo | Numbered, theorem head in Roman font and theorem text in italic style |
| thmstylethree | Numbered, theorem head in bold font and theorem text in roman style |

As per the output requirement, a corresponding new theorem style should be defined in the preamble area. For example, if you require "Proposition" environment to be set with "thmstyletwo", then you need to include the below lines in the preamble area:

```
\theoremstyle{thmstyletwo}
\newtheorem{proposition}{Proposition}
```

Refer to the "amsthm" package documentation for more details about the additional features available for new theorem styles. Apart from the above, a predefined "proof" environment is available. This environment prints a "Proof" head in italic font style and the "body text" in roman font style with an open square at the end of each proof environment.

```
\begin{proof}...\end{proof}
```

##### *7.6. Footnotes*

Footnotes are produced with the standard LaTeX command \footnote{<Some text>}. This typesets a numerical flag at the location of the footnote command and places the footnote text at the bottom of the page.

##### *7.7. Algorithms, Program codes, and Listings*

The "algorithm", "algorithmicx" and "algpseudocode" packages are used for setting algorithms in LaTeX. For algorithms, use the below format:

```
\begin{algorithm}
\caption{<alg-caption>}\label{<alg-label>}
\begin{algorithmic}[1]
...
\end{algorithmic}
\end{algorithm}
```

Refer to the above-listed package documentation for more details before setting algorithms.

For program codes, the "verbatim" package is required and the command to be used is \begin{verbatim}...\end{verbatim}.

The command `\begin{lstlisting}...\end{lstlisting}` is also used to set "verbatim" like environments. The listings package **supports highlighting of all the most common languages** and it is highly customizable. You will need to use the listings package for the {lstlisting} command to work, refer to the "lstlisting" package documentation for more details.

##### *7.8. Cross references*

A feature of LaTeX is the ability to automatically insert hypertext links within a document:

- the `\label{...}` command is used to identify an object (i.e. an equation, figure, table). This identifier is used in the `\ref{...}` command for cross-referencing.
- the `\ref{...}` command inserts a clickable link to an object as defined by it's label

##### *7.9. Citations*

The bibliography styles provided in the template package support author-year and numbered citations. The citation option used by default is the most common one for the bibliography style. Based on the preference of the journal you are submitting to, you can switch for 'sn-basic' and 'sn-chicago' between the options by adding or removing "Numbered" in the `\documentclass` line of the sn-article.tex file.

For example: `\documentclass[sn-basic,Numbered]{sn-jnl}`
For author-year citation style: `\documentclass[sn-basic]{sn-jnl}`

`\citep{...}` should be used for a parenthetical citation. For e.g. (Jones et al. 1990)
`\cite{...}` should be used for textual citation. For e.g. Jones et al. (1990)

Please refer to the "natbib" package documentation for full details and citation commands including optional arguments to describe chapters.

### 8. Back matter elements

##### *8.1. Heading levels in back matter*

Use the command `\bmhead{...}` for all the back matter heads which are to be placed below the `\backmatter` command.

##### *8.2. Appendices/Extended Data/Supplementary Information*

These sections are set with `\begin{appendices}...\end{appendices}` environment. All the other commands used to set section heads, tables, and figures inside this section remain the same as main body text.

##### *8.3. References*

BibTeX is the preferred format for references. BibTeX automates most of the work involved in references for articles. Using BibTeX options, both citations and references can be automatically updated to the preferred reference style. BibTeX works with two parts of the references: *content* and *style*. The *content* is stored separately in a plain text database file called .bib. The *style* and presentation of the .bib file content are processed with the help of BibTeX program using a style file called .bst (bibliography style file).

You are requested to use the sample bib file provided as a base for preparing your own .bib file. There are predefined bibliography style files available with this template. Select the bibliography style file according to the bibliography style as shown in the below table. Based on the preference of the journal you are submitting to, you can change the citation style (author-year or numbered) in the sn-article.tex file and the order of the reference list (alphabetical or numbered consecutively) in the bibliography style files. You will find more detailed instruction on how to do this in the respective files. For example, use "sn-nature" for Nature Portfolio journals.

| | **Bibliography style** | **Citation style** | **Option to be used** |
|---|---|---|---|
| **1** | Basic Springer Nature Nature/Chemistry Reference Style | authoryear | sn-basic |
| **2a** | Math and Physical Sciences Reference Style | numbered | sn-mathphys-num |
| **2b** | Math and Physical Sciences Reference Style | authoryear | sn-mathphys-ay |
| **3** | American Physical Society (APS) Reference Style | numbered | sn-aps |
| **4a** | Vancouver Reference Style | numbered | sn-vancouver-num |
| **4b** | Vancouver Reference Style | authoryear | sn-vancouver-ay |
| **5** | APA-based Social Sciences/Psychology Reference Style | authoryear | sn-apa |
| **6** | Chicago-based Humanities Reference Style | authoryear | sn-chicago |
| **7** | Nature Portfolio Reference Style | numbered | sn-nature |

For example, to use "Basic Springer Nature" reference style, you need to provide the option "sn-basic" to `\documentclass{...}` line as shown below: `\documentclass[sn-basic]{sn-jnl}`and extract the corresponding sn-basic.bst file from the bst folder and place it in the same folder as the sn-article.tex file.

Then include your .bib file at the end of your document as shown below:

\bibliography{<bib-file-without-extension>}

To generate a .bbl file run LaTeX on your manuscript file once, followed by BibTeX, and then run LaTeX twice, to ensure that references are set correctly.

The resulting bibliography is ready for typesetting with all formatting tags rendered according to the chosen reference style. For more details, please visit http://www.bibtex.org.

If you are submitting to one of the Nature Portfolio journals, using the eJP submission system, please include the references within the manuscript file itself. You may do this by copying the reference list from your .bbl file, and pasting it into the bibliography environment of the main manuscript .tex file. The basic bibliography environment 'thebibliography' must be used if you are using the natbib package.

For submissions to all other journals please check the submission guidelines on the respective journal website for information on the particular bibliography style that is being used for that journal.

### 9. Author support

Frequently asked questions are available at

https://www.springernature.com/gp/authors/campaigns/latex-author-support. If you cannot

find the answer you need in the FAQs, please contact.

### 10. Springer Nature Authoring Template – File details

| | | |
|---|---|---|
| User Manual | user-manual.pdf | Provides details on Springer Nature Authoring Template and its usage |
| Class file | sn-jnl.cls | Springer Nature authoring template class file |
| Sample LaTeX files | sn-article.tex | Sample for single column layout |
| Sample PDF file | sn-article.pdf | PDF output of sn-article.tex |
| .bst files | sn-basic.bst | .bst file to be used for Basic Springer Nature/Chemistry reference style |
| | sn-mathphys-num.bst sn-mathphys-ay.bst | .bst file (numbered/author year) to be used for Math and Physical Sciences Reference Style |
| | sn-aps.bst | .bst file to be used for American Physical Society (APS) Reference Style |
| | sn-vancouver-num.bst sn-vancouver-ay.bst | .bst file (numbered/author year) to be used for Vancouver Reference Style |
| | sn-apacite.bst | .bst file to be used for APA Reference Style |
| | sn-chicago.bst | .bst file to be used for Chicago-based Humanities Reference Style |

| | | |
|---|---|---|
| | sn-nature.bst | .bst file to be used for Nature Portfolio Reference Style |
| .bib files | sn-bibliography.bib | Common bib file |
| eps files | empty.eps | Dummy eps image used for placement only |
| | fig.eps | Dummy eps image used for placement only |

### 11. Change History

| **Version No.** | **Revision Date** | **Revision Note** |
|---|---|---|
| Version 1 | July 2021 | Original release date. |
| Version 2 | March 2023 | **April 2022**: Fixed Author affiliation bugs<br>**November 2021:** (a) .bst file switch; (b) sn-nature.bst is bringing in unnecessary information<br>**October 2021**: sn-mathphys.bst/line 652 error<br>**September 2021:** Avoid author names splitting over two lines |
| Version 2.1 | April 2023 | Author affiliation identifiers on the opening page -- fixed on the first compilation |
| Version 3 | December 2023 | sn-mathphys.bst has been split into Numbered and Author year style namely sn-mathphys-num.bst and sn-mathphys-ay.bst respectively<br><br>The Author Support contact email address has been updated in the User Manual.<br><br>An error in sn-aps.bst which did not output the title for the references when using inproceedings has been corrected. |
| Version 3.1 | December 2024 | FUNCTION {format.eprint} has been updated in sn-basic.bst, sn-nature.bst files to have linked urls.<br><br>amsthm package has been now called in the main class file sn-jnl.cls and removed from the sn-article.tex file<br><br>The display of commas has been corrected when using the \citep command.<br><br>sn-vancouver.bst has been split into Numbered and Author year style namely sn-vancouver-num.bst and sn-vancouver-ay.bst respectively<br><br>pdflatex option is now loaded as default |
